# Isolated loss of the AUTS2 long isoform, brain-wide or targeted to *Calbindin*-lineage cells, generates a specific suite of brain, behavioral and molecular pathologies

**DOI:** 10.1101/2023.05.04.539486

**Authors:** Yunshu Song, Christopher H. Seward, Chih-Ying Chen, Amber LeBlanc, Analise M. Leddy, Lisa Stubbs

## Abstract

Rearrangements within the *AUTS2* region are associated with a rare syndromic disorder with intellectual disability, developmental delay and behavioral abnormalities as core features. In addition, smaller regional variants are linked to wide range of neuropsychiatric disorders, underscoring the gene’s essential role in brain development. Like many essential neurodevelopmental genes, *AUTS2* is large and complex, generating distinct long (AUTS2-l) and short (AUTS2-s) protein isoforms from alternative promoters. Although evidence suggests unique isoform functions, the contributions of each isoform to specific *AUTS2-*linked phenotypes have not been clearly resolved. Furthermore, *Auts2* is widely expressed across the developing brain, but cell populations most central to disease presentation have not been determined. In this study, we focused on the specific roles of AUTS2-l in brain development, behavior, and postnatal brain gene expression, showing that brain-wide AUTS2-l ablation leads to specific subsets of the recessive pathologies associated with C-terminal mutations that disrupt both isoforms. We identify downstream genes that could explain expressed phenotypes including hundreds of putative direct AUTS2- l target genes. Furthermore, in contrast to C-terminal *Auts2* mutations which lead to dominant hypoactivity, AUTS2-l loss-of-function is associated with dominant hyperactivity, a phenotype exhibited by many human patients. Finally, we show that AUTS2-l ablation in *Calbindin 1*-expressing cell lineages is sufficient to yield learning/memory deficits and hyperactivity with abnormal dentate gyrus granule cell maturation, but not other phenotypic effects. These data provide new clues to *in vivo* AUTS2-l functions and novel information relevant to genotype-phenotype correlations in the human *AUTS2* region.

## Introduction

*AUTS2* was first discovered as the gene disrupted by translocation in a pair of Autistic twins (Sultana et al., 2002), and since that initial finding, mutations in this gene have been associated with a remarkably wide variety of neurological and developmental disorders. Individuals carrying genomic rearrangements in the *AUTS2* region typically express “AUTS2 syndrome” phenotypes, typified by short stature, distinctive facial features, developmental delay and intellectual disability (ID), with some combination of autism spectrum disorders (ASD), feeding difficulties, seizures, hyperactivity and other features (Beunders et al., 2015, 2016; Sanchez-Jimeno et al., 2021). More recently, in-frame deletions within exon 9 that ablate the essential HX domain of the protein were associated with Rubenstein-Taybi syndrome (RTS) (Liu et al., 2021); a missense mutation within the HX domain has also been reported in an individual with profound ID, epilepsy, and brain pathology including microcephaly (Fair et al., 2023). In addition to these mutations, single nucleotide polymorphisms (SNPs) throughout the region are associated with attention deficit hyperactive disorder (ADHD) (Beunders et al., 2015; Amarillo et al., 2014; Elia et al., 2009), schizophrenia (Zhang et al., 2014), dyslexia (Amarillo et al., 2014; Palumbo et al., 2021), and susceptibility to substance abuse (Chen et al., 2013; Schumann et al., 2011). The association of this single genetic region with such a wide variety of neurodevelopmental disorders suggests a central and widespread role in brain development.

Like many essential neurodevelopmental genes, *AUTS2* is large and complex, spanning more than 1.5 Mb and encoding long (AUTS2-l) and short (AUTS2-s) protein isoforms from conserved alternative promoters (Beunders et al., 2015; Geng et al., 2022; Monderer-Rothkoff et al., 2021; Weisner et al., 2019). Analysis of mouse mutations and neurons *in vitro* have suggested a wide variety of molecular functions including regulation of the actin cytoskeleton (Hori et al., 2014) and ubiquitin-mediated protein degradation (Geng et al., 2022), as well as nuclear functions including the regulation of neurodevelopmental gene expression through chromatin remodeling (Gao et al., 2014; J. Li et al., 2022a; S. Liu et al., 2021) and RNA stabilization (Castanza et al., 2021). Studies with embryonic stem cell (ESC)-derived neurons (Monderer-Rothkoff et al., 2021; Liu et al., 2021) and brain organoids (Allegra et al., 2017; Fair et al., 2023) have shown that *Auts2* is involved in basic steps of neurogenesis and neuron maturation, and combined *in vivo* and *in vitro* studies have pointed to a special function in synapse formation the regulation of dendritic spines (Hori et al., 2020). These studies have provided intriguing hints that the functions of AUTS2-l and AUTS2-s proteins are distinct, but likely intertwined. For example, AUTS2-s is expressed at early stages of neuron commitment *in vitro,* whereas AUTS2-l rises later as the cultured neurons begin to mature (Monderer-Rothkoff et al., 2021). At the molecular level, AUTS2-l is the only isoform that associates with PRC1 complexes (Gao et al., 2014; S. Liu et al., 2021; Monderer-Rothkoff et al., 2021), while AUTS2-s, which can also function as a transcriptional activator (Monderer-Rothkoff et al., 2021), must operate via alternative mechanisms. Some evidence suggests that the two isoforms might act additively or cooperatively; for example, human mutations in exons 1-7, which are uniquely coding in AUTS2-l, are associated with many of the AUTS2 syndrome phenotypes although with mild expression, while mutations in exons encoding C-terminal amino acids that are shared by AUTS2l- and AUTS2-s (exons 8-19) generate more severe forms of syndromic disease. Consistently, animals homozygous for brain-wide conditional knockout (cKO) of *Auts2* exon 7 are born live with subtle developmental phenotypes (Gao et al., 2014), whereas mice homozygous for germline or brain-wide C-terminal mutations die perinatally with significant brain pathologies (Castanza et al., 2021; Hori et al., 2014).

Therefore, *Auts2* is clearly required for normal brain development and its malfunction or dysregulation can lead to neurodevelopmental disease. However, the contributions of AUTS2 protein isoforms to specific aspects of disease presentation have not been clearly elucidated. The distinct and intertwined functions of AUTS2 isoforms must be clarified before mechanisms of AUTS2-linked neurological disorders can be understood and before genotype: phenotype correlations in this essential genomic region can be resolved. In this study, we used the published *Auts2* exon 7 cKO mutant (Gao et al., 2014) to address several open questions about the developmental functions of AUTS2-l. For example, although AUTS2-l appears to be the dominant isoform in cerebellum (Weisner et al., 2019; Yamashiro et al., 2020) the impact of isolated loss of AUTS2-l function on cerebellar development has not been investigated. Additionally, although recessive phenotypes have been described for animals inheriting cKO of *Auts2* exon 6 (J. Li et al., 2022a), AUTS2-l loss-of-function (LOF) in that study was driven specifically in cortical excitatory neurons, although *AUTS2* is expressed more broadly across the developing brain (Bedogni et al., 2010; Weisner et al., 2019; Yamashiro et al., 2020), and potential dominant phenotypes have not been examined. We designed experiments to address these questions with the goal of clarifying functions of AUTS2-l in the developing brain, and to leverage these data to suggest *in vivo* functions for AUTS2-s as well.

## Materials and Methods

### Animals and Tissue Collection

The *Auts2*-ex7^fl/fl^ allele (*Auts2^tm1.1Dare^*; Gao et al., 2014), was provided as a kind gift from the laboratory of Dr. Danny Weinberg (NYU). *Nes-cre* (*B6.Cg-Tg(Nes-cre*)1Kln/J), *Calb1-cre* (*B6;129S- Calb1tm2.1(cre)Hze/J*) and *Cmv-cre* (*B6.C-Tg(CMV-cre)1Cgn/J*) were purchased from the Jackson Laboratory. *Cre*-expressing strains were crossed to *Auts2*-ex7^fl/fl^ animals to generate homozygotes, heterozygotes and wild-type littermates for behavioral testing and tissue collection. WT, *Nes-cre*, *ex7^-/-^*and heterozygote germline KO *Cmv-cre*, *ex7^-/+^* were tested for developmental milestones including body weight, righting reflex, eye opening, open field test, NOR, and sociability and social memory tests as previously described (Weisner et al., 2019; Chen et al., 2022). This study strictly followed the Guide for the Care and Use of Laboratory Animals of the Use Committee of the University of Illinois (Animal Assurance Number: A3118-01; approved IACUC protocol number 18240) and the Pacific Northwest Research Institute (A3357-01; IACUC protocol 201-20). Animals were maintained under standard conditions (12h light/dark cycle, group housed); all tests were done between 1-5hr prior to lights-off or 1- 5hr prior to lights-on.

### Behavioral Tests

#### Righting reflex

righting reflex test was modified from previous study (Castelhano-Carlos et al., 2010; Chen et al., 2022) with following modifications. On postnatal day 5, pups were placed belly up on flat surface and scored based on time lapsed before they get back to their feet on the ground: score 0 = failed to turn back within 15 sec; score 1 = pups turned between 10-15 sec; score 2 = pups turned within 10 sec.

#### Eye opening

Eye opening was measured on postnatal day 13 following previous study (Chen et al., 2022). In brief, scores from 0-2 are assigned. Score 0 = both eyes are closed; score 1 = one eye is open or both eyes partially open; score 2 = both eyes are fully open.

#### Open field test

mouse was placed in a novel open arena, 35cm x 30xm paper box for 10min and videotaped for analysis using DeepLabCut 2.2.3 (DLC) (Mathis et al., 2018). Tracking framework was trained to recognize tail base, middle back, nose tip, left ear and right ear of the mouse. A skeleton, lines connecting between points, was included to connect the tail base to middle back, middle back to nose tip, and both ears each to the nose tip. We used default neural network and iteration settings. Videos are cropped using GUI to include only the arena. Middle back point coordinates were used to calculate total travel distance in pixels from 18,000 frames (10 mins).

#### Stereotypic behavior (grooming)

The average grooming time per episode during the open field test was measured following previously described protocols (Chen et al., 2022).

#### Novel object recognition (NOR)

Published protocols were followed (Botton et al., 2010; Chen et al., 2022). In brief, tasks include three sessions: habituation, training and retention. During habituation, mouse completed open field test for 10min. After 24hrs interval, same mouse was introduced to the same arena with two novel objects for training session, and mouse was allowed to explore the objects for 10mins. The total time a mouse was actively exploring the objects was quantified as total exploration time for NOR- exploratory activity. 24 hours later, during retention session, one of the familiar objects that appeared in training session was exchanged for a new object in the same arena, and the mouse was allowed to explore for 5 min. The exploratory index is calculated as percentage of time mouse spent in exploring novel object/ total time spent exploring both objects in retention session. Exploratory behavior was defined as sniffing or touching the object.

#### Three chamber test

we followed the protocols described in published studies (Chen et al., 2022; Hori et al., 2015) with minor modifications. A Plexiglass box was divided into 3 chambers: object chamber, middle chamber, and social chamber. The tested mouse was introduced to the empty box and allowed to explore for 10mins. Then a novel object (Lego toy or items of similar size/complexity) was introduced to the object chamber, and a sexually immature juvenile female mouse (P21-P28) was placed in the social chamber as stranger mouse 1. Both the object and the stranger mouse were restrained under a small stainless steel wire cup. The tested mouse was again allowed to explore the box for another 10mins. Time of the mouse spent in each chamber was quantified, and the percentage of time spent in the social chamber/time spent in both object chamber and social chamber was quantified for sociability behavior. Lastly, a novel sexually immature juvenile female mouse (P21-P28) was introduced to replace the object as stranger mouse 2, and tested mouse was allowed to explore the box now with a familiar mouse (stranger mouse 1) and a novel mouse (stranger mouse 2) freely for 10min. Social memory is quantified by percentage of time the tested subject spent in novel mouse chamber / total time spent in either familiar mouse chamber or novel mouse chamber during the final 10min session.

### RNA collection, RNA-seq, qPCR, and RiboTagging

#### RNA, qRT-PCR

RNA and cDNA were prepared as previously described (Weisner et al., 2019). qRT-PCR was done using custom-designed primers for specific *Auts2* isoforms (**Supp. Table 1**), and expression values were normalized relative to the *Pgk1* control in each sample and compared using standard methods (Livak & Schmittgen, 2001).

**Table 1.** Developmental and behavioral phenotypes in ex7 mutants compared to other published.

FIGURE LEGENDS
| Phenotype | *Nes-cre*<br>ex7<sup>-/-</sup> | *Cmv-cre*<br>ex7<sup>+/-</sup> | *Calb1-cre*<br>ex7<sup>-/-</sup> | *Emx1-cre*<br>ex6<sup>-/-a</sup> | *Auts2*<br>del8/+<sup>b</sup> | *Auts2*<br>neo/+<sup>c</sup> | 16Gso/<br>16Gso<sup>d</sup> |
| --- | --- | --- | --- | --- | --- | --- | --- |
| Perinatal survival | N | N | N | N | N | N | N |
| Postnatal growth | - | - | N | N | - | - | - |
| Developmental milestones<sup>e</sup> | - | - | - | *nr* | *nr* | *nr* | - |
| Stereotypic behaviors | N | + | N | + | *nr* | *nr* | + |
| Travel distance, open field | + | + | + | N | - | - | N<sup>e</sup> |
| Exploratory activity | + | N | N | *nr* | *nr* | N | - |
| Cognitive memory (NOR) | - | N | - | N | - | - | - |
| Sociability | - | N | N | N | - | N | - |
| Social memory | N | N | *nt* | - | - | N | *nt* |

+ = increased; - = decreased; N = normal; *nr* = not reported; *nt* = not tested
<sup>a</sup> Li *et al.*, 2022
<sup>b</sup> *CAG-cre*, ex8<sup>+/-</sup>; Hori *et al.*, 2020.
<sup>c</sup> Hori *et al.*, 2015; homozygotes for this genotype do not survive.
<sup>d</sup> Weisner, Chen *et al.*, 2019; activity levels were normal in homozygotes, but significantly increased in heterozygotes measured as home cage activity.
<sup>e</sup> timing of righting reflex development and/or eye opening

#### RNA-seq

Three P0 *Cmv-cre*, *Auts2*-ex7^-/-^ and WT forebrains, combined midbrain and hindbrains together, and P14 *Nes-cre*, *Auts2*-ex7^-/-^ HC RNA samples were prepared as previously described (Weisner et al., 2019), and RNA-seq libraries were generated and analyzed as previously described (Chen et al., 2022). Briefly, libraries were constructed with Illumina TruSeq RNA library Prep Kits v2. and sequenced on Illumina Hi-Seq 4000 sequencer for 150bp paired end reads (17-30M reads/sample) through Genewiz sequencing service (South Plainfield, NJ). STAR v2.7.5a was used to map reads to mm10/GRCm38 genome and ensemble v102 annotation). Differential expression analysis was done using EdgeR v3 (Robinson et al., 2009) as described in Saul and Seward et al. (Saul et al., 2017) using Benjamani-Hochberg correction to assign fdr values to particular genes. Genes with fdr < 0.05 were considered to be significantly differentially expressed. For functional analysis, we used ToppCluster annotation tool (Kaimal et al., 2010) with default statistical parameters, focusing on genes identified as having at least 1.5-fold change (FC) in mutants compared to WT controls with FDR less than 0.05. Visualization of RNA-seq data was performed with “ggplot2” package(Wickham H, 2016).

#### Ribo-Tag immunoprecipitation

*Calb1-cre* (*B6;129S-Calb1tm2.1(cre)Hze/J*) mice were crossed to *Rpl22-*HA strain (B6N.129-Rpl22tm1.1Psam/J) (Sanz et al., 2009) from Jackson Laboratories. Immunoprecipitation (IP) enrichment was performed on RNA isolated from dissected brain tissue as described in by Sanz et al. (Sanz et al., 2009). In brief, samples were suspended in homogenization buffer (50mM Tris, pH7.5, 100mM KCl, 12mM MgCl2, 1%NP40-substititue, 1mM DTT, 200U/mL Promega RNasin, 1mg/mL heparin, 100μg/mL cycloheximide, 2x PIC (Roche)). HA-antibody (ab9110, Abcam) was added to the supernatant after 10,000g x 10 min centrifugation at 4C, and incubated at 4C with rotation for 4hr. Then protein G beads pre-equilibrated with homogenization buffer were added to the supernatant-antibody mix to bind overnight at 4C. Beads were washed with high salt buffer (50mM Tris, pH7/5, 300mM KCl, 12mM MgCl2, 1% NP40 substitute, 1mM DTT, 100μg/mL cycloheximide) three times, and 900μL of trizol was added to the beads to release RNA from the HA-labeled polysomes.

### Immunohistochemistry and histopathology

Isolated tissues were fixed in fresh 4% paraformaldehyde, embedded in paraffin, and cut into 5 micron sections using a Leica RM2155 microtome and Super Plus charged slides (Leica). Cerebellum (CB) sections were sectioned sagittally; hippocampus (HC) was sectioned coronally. For histology experiments, deparaffinization, rehydration and antigen retrieval steps were followed and paraffin-embedded slides were stained with hematoxylin and eosin (H&E) as previously described (Weisner et al., 2019). IHC blocking was done in Antibody Diluent Reagent Solution (Life Technologies), and primary antibodies were tested and used at the optimal ratio in the same antibody diluent and incubated overnight at 4C. Secondary antibodies (1:200) were also diluted in the same antibody diluent for 1hr at room temperature. Hoechst 3342 was used for nuclear stain prior to scanning with Leica Sp8 confocal microscope and LAS imaging system. Primary antibodies: AUTS2 (Sigma, HPA000390), Calbindin D- 28K (Sigma C9848, 1:300), Calbindin (Santa Cruz sc-365360, 1:300), Calretinin (Santa Cruz sc-365989, 1:300), Calretinin (ZYM MD, 1:300), Tbr2 (Invitrogen 14-4875-52, 1:100), Parvalbumin (R&D system AF5058, 1:50 – 1:100), cFOS (Santa Cuz s-166940, 1:200), GAD (R&D system AF2086-SP, 1:100); Secondary antibodies (Thermo-Fisher Scientific): Goat anti-mouse IgG (Alexa Fluor 488, A11001), Goat anti-Rabbit IgG (Alexa Fluor 594, A11012), Donkey anti-Rat IgG (Alexa Fluor 594, A21209), Donkey anti-Sheep IgG (Alexa Fluor 594, abcam, ab150180), Donkey anti-Sheep IgG (Novus Biologicals, DyLight488, NBP173002). All measurements and counts were done on multiple sections from at least N=3 animals for each genotype.

#### Cell number counts, structure thickness and analysis

The number of cells stacked in vertical columns of the dentate gyrus (DG) granule cell layer (GCL) structure was counted. The thickness of both CalB+ and CalR+ layers was measured and the number of cells / column of CalB+ and CalR+ cells were counted. The ratio of CalB+/CalR+ layer or cell counts were reported. For cell density counts, the thickness of each structure (DG GCL or *Cornu Ammonis* (CA) pyramidal cell layer (PCL) was measured, and the number of cells inside a square, with side-length equal to the thickness of structure, were counted. Cell density was calculated by dividing total cell number counted by the squared area. Analysis performed with NanoZoomer NDP view Digital Pathology software. Two or more measurements were taken per section on 2-6 sections from at least N=3 animals of each genotype. Structural thickness was determined from two or more measurements per section on 2-9 sections from at least N=3 animals of each genotype. Suprapyramidal projection bundle thickness and EGL thickness were determined by averaging three measurements from each section on 3-11 sections from at least N=3 animals of each genotype. Statistical significance was determined through two-tailed t-test. P-value ranges are reported for each figure respectively. Statistical significance was determined through two-tailed t-test. P-value ranges are reported for each figure respectively.

### Statistical analysis

All statistical analysis and graph presentation are performed with GraphPad Prism 8 (GraphPad Software, San Diego, California USA, www.graphpad.com) unless specially noted. Two-tailed unpaired t-tests were used to determine statistical significance for behavioral tests, qRT-PCR, and cell counts and outliers were excluded based on specified outlier tests reported in figure legends. P values less than 0.05 were considered significant; ranges and other details are indicated in figure and figure legends. Number used for each test are reported in Methods, with further information in the text, figure legends, or in the Supplemental Tables in context with discussion of the data. Graphs are presented as mean ± standard error of the mean (SEM). Statistical analysis for RNA-seq data is detailed above.

## RESULTS

### Conditional deletion of *Auts2* exon 7 leads to AUTS2-l ablation, but transient up-regulation of intact AUTS2-s transcripts and protein in neonatal brain

Animals in which *Auts2* exon 7 has been deleted homozygously with the brain-wide *Nestin cre* allele (hereafter abbreviated as *Nes-cre*, ex7^-/-^ animals) are viable, and were reported to show an absence of AUTS2-l protein in brains with expression of an intact AUTS2-s protein (Gao et al., 2014). In addition to *Nes-cre*, ex7^-/-^ mice, we generated *Cmv-cre*, ex7^-/-^ animals which stably inherit a germline deletion of *Auts2* exon 7; this latter genotype had not been examined previously. Like the *Nes-cre,* ex7^-/-^ mice (Gao et al., 2014), *Cmv-cre*, ex7^-/-^ homozygotes were viable but smaller than WT littermates (**Table 1**). As reported, *Nes-cre,* ex7^-/-^ mice were long-lived and otherwise healthy although they were sterile. In contrast, some but not all *Cmv-cre*, ex7^-/-^ mice survived into late adulthood. Heterozygotes of both genotypes were healthy and long-lived.

First, we collected RNA and protein from mutants to confirm and measure AUTS2 isoform expression. We tested forebrain (FB) from newborn (postnatal day 0.5, hereafter abbreviated P0) *Cmv-cre*, ex7^-/-^ and WT littermates and combined midbrain and hindbrain (MH) regions from the same animals. We also tested RNA and protein from dissected hippocampus (HC) and cerebellum (CB) of *Nes-cre*, ex7^-/-^ mice at P14, a critical stage for development of both HC and CB (Berg et al., 2019; Brandt et al., 2003; Nicola et al., 2015; Sudarov et al., 2007). We first confirmed exon 7 deletion in mutant *Auts2* transcripts using reverse transcript PCR (RT-PCR) to generate cDNA fragments from the P0 samples and examining PCR product size (**Fig. 1a**) followed by Sanger sequencing (not shown). We then examined the levels of each transcript with quantitative RT-PCR (qRT-PCR) (**Table S1**). Although exon 7 deletion clearly puts AUTS2-l transcripts out of frame, mutant transcript levels were not significantly different from WT in either P0 FB or MH samples (**Fig 1b, left panel**); this result suggested inefficient clearance by nonsense mediated decay in the neonatal brains. In contrast, AUTS2-l transcript levels were significantly reduced in P14 HC (**Fig. 1b, right panel**). However, consistent with published reports (Gao et al., 2014; S. Liu et al., 2021) AUTS2-l protein was ablated in ex7^-/-^ mutant brains (**Fig 1c, left panel, red arrow**).

**Figure 1.**
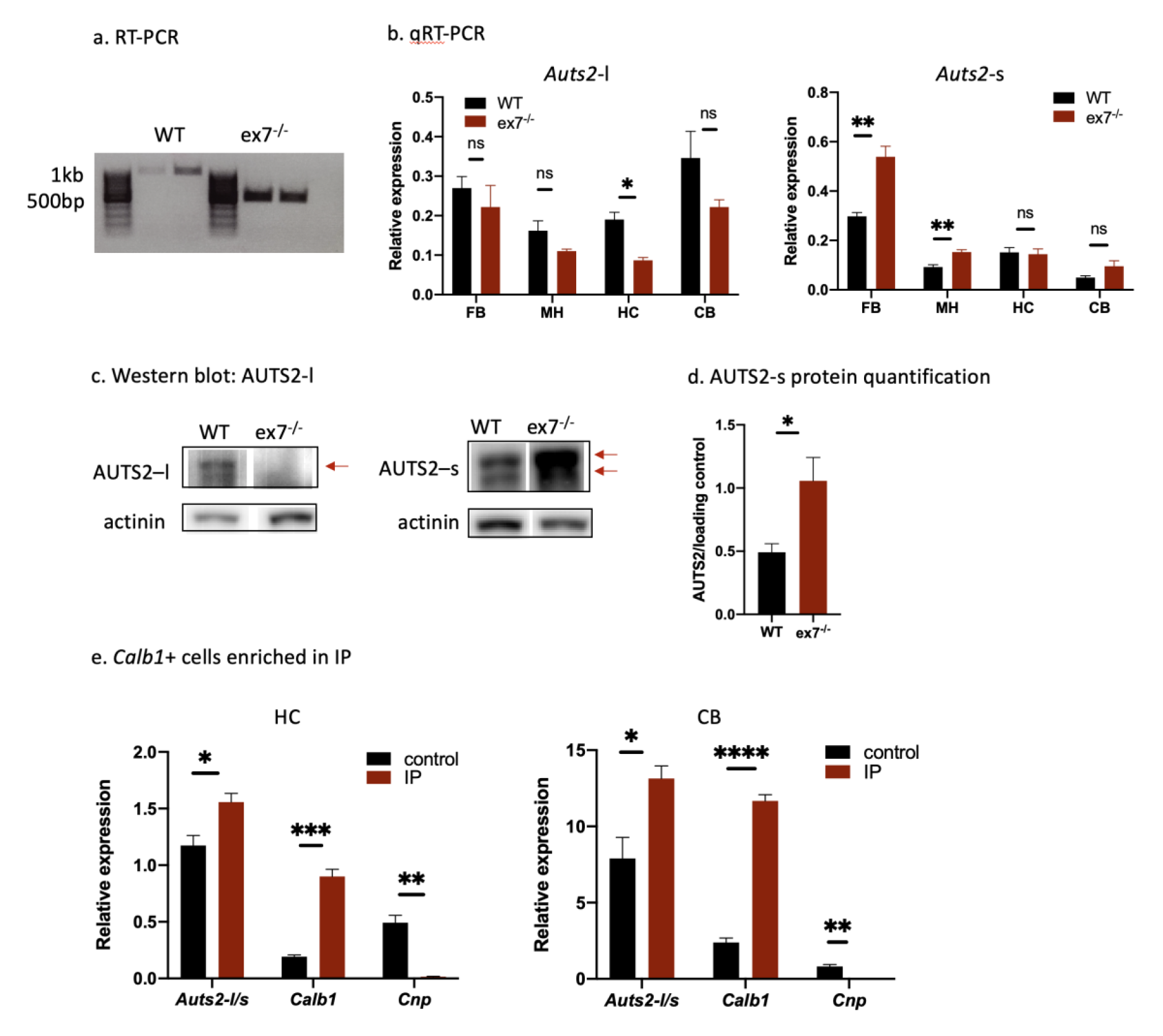
**Expression of *Auts2* isoforms in mutant and wild type brain**. **a)** Confirmation of exon 7 deletion in *Cmv-cre,* ex7^-/-^ brains. Reverse transcript (RT)-PCR products from brains of WT mice are ∼500bp longer than those from ex7^-/-^ mice, confirming the deletion of the 470bp exon 7 as expected. **b) AUTS2-l and AUTS2-s transcript expression** characterization by qRT-PCR in RNA from WT and ex7^-/-^ animals. RNA from P0 *Cmv-cre,* ex7^-/-^ forebrain (FB) and combined midbrain-hindbrain (MH), or from P14 *Nes-cre,* ex7^-/-^ hippocampus (HC) and cerebellum (CB) was analyzed. **c) Representative western blot** analysis showing AUTS2-l (left) and major AUTS2-s (right) protein isoform expression in WT and ex7^-/-^ animals and the actinin loading control. Red arrows point to AUTS2 bands. **d) quantification of AUTS2-s** band intensity from western blot in panel (c), using FIJI-ImageJ. **e) Ratio of AUTS2-l/AUTS2- s transcripts**, and relative expression of *Calb1* and *Cnp* transcripts in HC (left) and CB-right in whole HC (control) and immunoprecipitated (IP) RNA from CalB+ cells in *Calb1-cre,* RiboTag mice. Two-tailed t-test was used to test statistical significance, * = p<0.05, ** = p<0.01, **** = p<0.0001, ns = not significant.

Testing next for expression of AUTS2-s, we were surprised to find that the transcripts were significantly *up-regulated* in both P0 FB and MH fractions (**Fig. 1b, right panel**). Furthermore, the AUTS2-s protein, which appeared as a doublet (probably due to the documented alternative splicing of exon 10) around 100 kD in size, was also significantly more abundant in the P0 mutant than WT brain (**Fig. 1c right panel, red arrows, 1d)**. In contrast to P0, at P14 AUTS2-s transcripts and protein were expressed at WT levels in mutant hippocampus; AUTS2-s transcript levels trended higher in the P14 mutant cerebellum, although the differences did not reach statistical significance (**Fig. 1b, right panel**). The data suggested a possible feedback loop in which AUTS2-l regulates expression of AUTS2-s, either directly or indirectly but transiently, in neonatal brain.

### AUTS2-l LOF has subtle impact on cerebellar development, but dramatic hippocampal effects

Specific pathologies in CB (Yamashiro et al., 2020) and HC (Castanza et al., 2021; Hori et al., 2020; Weisner et al., 2019) have been reported for mice homozygous for C-terminal *Auts2* alleles. Some of the same hippocampal phenotypes were also recently reported for *Emx1-cre*, ex6^-/-^ mice, tying these pathologies to AUTS2-l LOF in forebrain excitatory cells (Li et al., 2022) (**Table 1**). Our next goal was to investigate pathologies associated with the brain-wide ablation of AUTS2-l, focusing on HC and CB in *Nes-cre*, ex7^-/-^ mice at P14 and in adults (P60) (**Fig. 2,3**).

**Figure 2.**
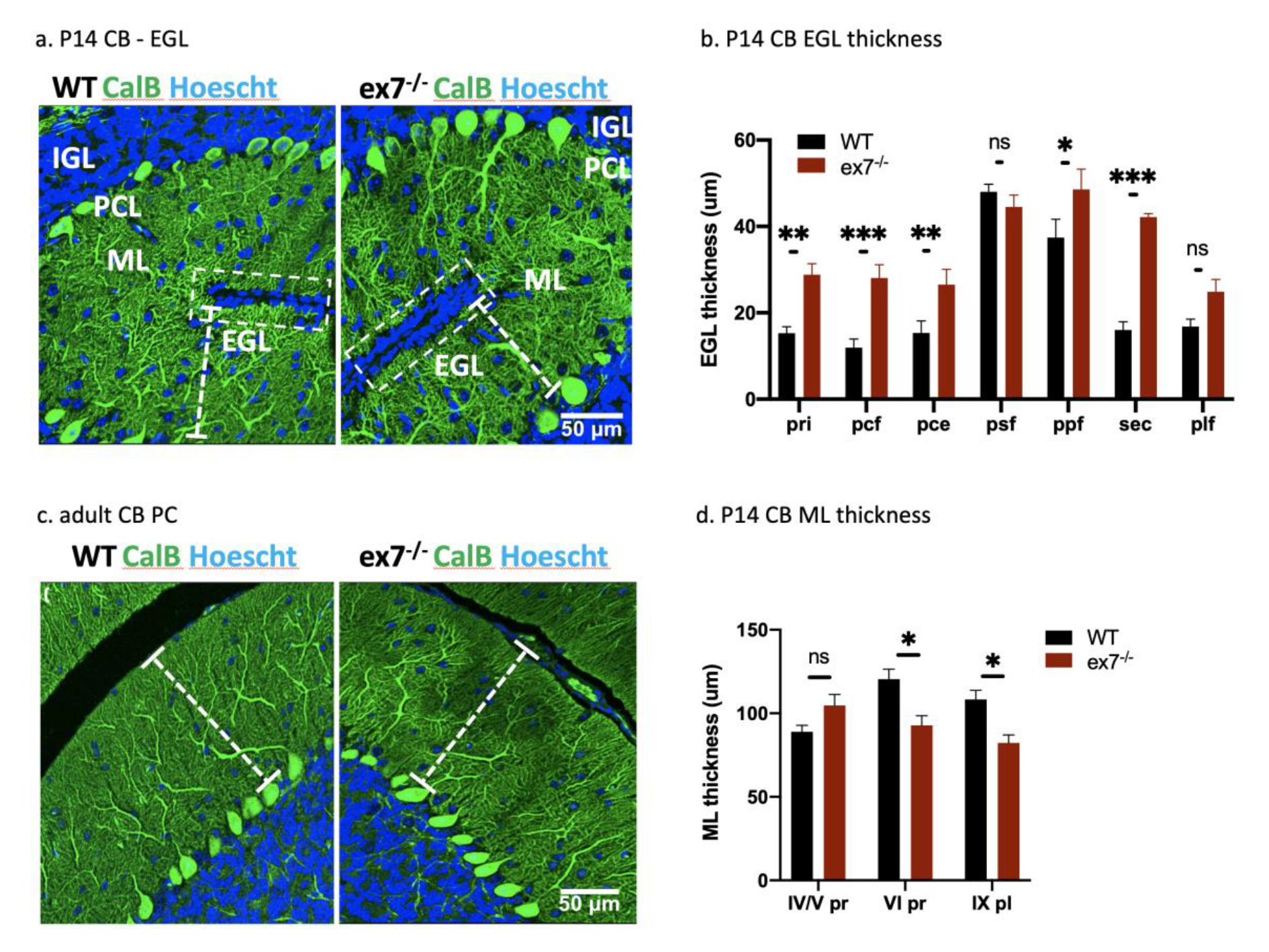
**Cerebellar pathology in *Nes-cre,* ex7^-/-^ mutant mice**, abbreviated simply as ex7^-/-^ in this figure. **a) Representative image of the base of foliation in P14 WT and *Nes-cre*, ex7^-/-^ cerebellum**; white boxes enclose the external granule layer (EGL); white dashed lines show the thickness of the molecular layer (ML); Blue =Hoechst, green = CalB; PCL = Purkinje cell layer. **b) quantification of EGL thickness** in different fissures in P14 WT and *Nes-cre*, ex7^-/-^ cerebellum. WT n=3-5, mutant n=5-6 sections were measured from 3 animals each genotype. pri = primary fissure, pcf = preculminate fissure, pce = precentral fissure, psf = posterior superior fissure, ppf = prepyramidal fissure, sec = secondary fissure, plf = posterolateral fissure; **c) Representative image of PC branching** in adult WT and *Nes-cre*, ex7^-/-^ mice; white dashed lines measure thickness of ML; **d) measurements of ML thickness** in P14 WT and *Nes-cre*, ex7^-/-^ vermis. Two-tailed t-test was used to test statistical significance, * = p<0.05, ** = p<0.01, **** = p<0.0001, ns = not significant.

**Figure 3.**
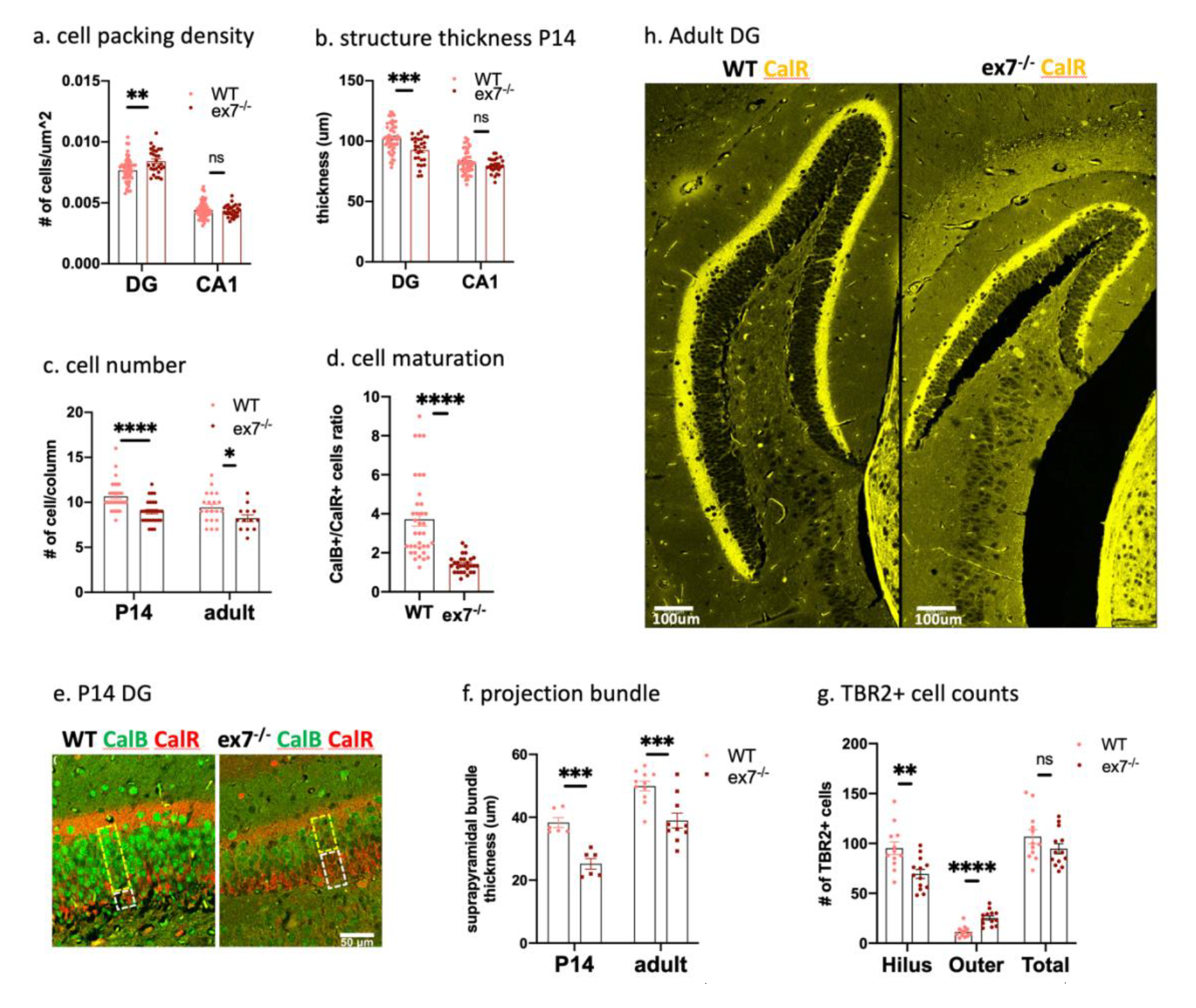
**Hippocampal (HC) pathology in *Nes-cre,* ex7^-/-^ mutants,** abbreviated as ex7^-/-^ in figures. **a) Dentate gyrus (DG) granule cells** were more densely packed in P14 *Nes-cre*, ex7^-/-^ HC as compared to WT, but CA1 pyramidal cells were normally packed **b) DG structure thickness** was significantly decreased in P14 *Nes-cre*, ex7^-/-^ mice compared to WT animals, although CA1 structure thickness was not significantly different in WT and mutant mice. **c) The total number of cells per column** in the DG was significantly reduced in both P14 and adult *Nes-cre*, ex7^-/-^ HC compared to WT littermates. **d) The ratio of CalB+/CalR+ cells** was significantly decreased in *Nes-cre*, ex7^-/-^ mutants DG compared to WT littermates. **e) Representative picture of the CalB+ and CalR+ layers** in P14 WT and *Nes-cre*, ex7^-/-^ HC. The yellow box highlights the CalB+ cell layer and the white box shows CalR+ cell layer in each sample. **f) The thickness of the suprapyramidal bundle** of the DG gc-CA3 projection was significantly reduced in P14 and adult *Nes-cre*, ex7^-/-^ animals compared to WT HC. **g) The number of TBR2+ cells** was significantly reduced in the hilus and DG gc layer (Hilus column) and increased in inner molecular layer (IML)(Outer column) of the DG in P14 *Nes-cre*, ex7^-/-^ HC compared to WT littermates, although the total number of TBR2+ cells was not changed. **h) HMN collateral axon tracks** stained with CalR show a bright signal and normal thickness in adult *Nes-cre*, ex7^-/-^ HC. Two-tailed t-test was used to test statistical significance: * = p<0.05, ** = p<0.01, ***= p<0.001, **** = p<0.0001, ns = not significant.

First, we tested the expression of AUTS2-s and AUTS2-l transcripts in two major cell types with documented *Auts2* mutant pathology: cerebellar Purkinje cells (PC) and maturing post-mitotic hippocampal dentate gyrus granule cells (DG gc). Our previous data suggest that AUTS2-l is the major isoform expressed in P14 CB whereas AUTS2-s is dominant in neonatal forebrain including P14 HC, although AUTS2-l is expressed in P14 HC as well (Castanza et al., 2021; Liu et al., 2021; Weisner et al., 2019). Since the Calbindin-D28K protein (abbreviated CalB, encoded by gene *Calb1*) is a well-established marker for both PC and maturing DG gc (Barski et al., 2003; Li et al., 2017), we used the *Calb1-cre* (Daigle et al., 2018) in combination with the RiboTag allele (Sanz et al., 2009) to isolate PC-or DG gc-enriched ribosome-associated RNA from WT P14 CB and HC. We then used qRT-PCR to compare the RiboTagged RNA to unselected total RNA from each brain region. Testing the enrichment of transcripts from CalB-expressing (CalB+) cells, we confirmed the robust enrichment of *Calb1* transcripts in the RibTagged RNA, as expected, along with depletion of *Cnp,* a marker of oligodendrocytes (**Fig. 1e**).

Next, we tested for relative levels of AUTS2-l and AUTS2-s transcripts in the HC samples and found that although AUTS2-s was indeed in highest abundance overall, the AUTS2-l/AUTS2-s ratio was significantly higher in RiboTag-enriched compared to the unselected RNA (**Fig 1e, left panel**). In CB, AUTS2-l transcripts dominated the unselected samples, and were further enriched in the RiboTagged RNA (**Fig. 1e, right panel**). The data suggested that in both HC and CB, AUTS2-l transcripts are expressed and relatively enriched in cells of the CalB+ lineage, indicating that AUTS2-l could play an important role in phenotypes expressed in those cells.

With that information in mind, we next examined *Nes-cre,* ex7^-/-^ mutant brains for pathologies that have been documented for other *Auts2* alleles. Given that AUTS2-l is the dominant isoform in both cerebellum overall and PC specifically at P14, it seemed logical to assume that the depletion of AUTS2-l would be associated with the dramatic cerebellar phenotypes described for C-terminal mutations. However, although we observed subtle abnormalities in the *Nes-cre,* ex7^-/-^ cerebellum we did not see the expected dramatic effects. Specifically, we found that the external granule layer (EGL) was significantly thicker in mutants than in WT littermates at P14, suggesting a delay in granule neuron migration from EGL to form the inner granule layer (Sudarov et al., 2007) (**Fig. 2a-b**). In contrast, the P14 mutant CB molecular layer was decreased in thickness compared to WT littermates (**Fig. 2a,d**), suggesting insufficient PC dendritic branching at this stage. Certain cerebellar lobules were more severely affected than others; for example, in the vermis, lobules VI pr, IX pl showed significantly decreased molecular layer thickness while lobules IV/V pr did not (**Fig. 2d**). However, the decreased thickness observed at P14 did not persist in those same lobules into adulthood (**Supp. Fig. 1a**). This finding could suggest a delay in PC dendritic growth, although we did not detect any obvious abnormalities in overall dendritic branching or structure of PCs in *Nes-cre*, ex7^-/-^ adults (**Fig. 2c**). Previously reported phenotypes including the loss of PCs with age, misalignment of the PC layer (Weisner et al., 2019), and obviously and significantly stunted PC dendrites in adults (Weisner et al., 2019; Yamashiro et al., 2020) were also not observed (**Table 2**). Therefore, despite the dominant expression of AUTS2-l in developing postnatal PC, AUTS2-l LOF is not sufficient to recreate the dramatic defects in PC maturation and survival.

**Table 2.** Pathology observed in hippocampus (HC) or cerebellum (CB) in *Auts2*-ex7 mutations compared with those identified in published *Auts2* mouse mutants.

|  | *Nes-cre*,<br>ex7<sup>-/-</sup> | *Calb1-cre*,<br>ex7<sup>-/-</sup> | *Emx1-cre*,<br>ex6<sup>-/-a</sup> | ex8<sup>-/-b</sup> | *Emx1-cre*,<br>ex15<sup>-/-c</sup> | 16Gso/<br>16Gso<sup>d</sup> |
| --- | --- | --- | --- | --- | --- | --- |
| HC DG – overall size | - | N | - | *nr* | - | - |
| HC DG –mature/immature gc ratio | - | - | N | *nr* | *nr* | - |
| HC DG – TBR2+ INP numbers | N | N | - | *nr* | - | + |
| HC DG - ectopic TBR2+ INP | + | N | + | *nr* | *nr* | *nr* |
| HC suprapyramidal bundle thickness | - | *nt* | - | *nr* | *nr* | *nr* |
| HC CA1- abnormal excitation (c-Fos) | + | N | + | + | + | + |
| HC PV+ interneuron cell number | - | N | *nr* | *nr* | *nt* | *nr* |
| HC HMN collateral axon track | N | N | *nr* | *nr* | - | N<sup>e</sup> |
| CB PC numbers/survival | N | *nt* | *nr* | - | *nr* | - |
| CB P14 EGL thickness | + | *nt* | *nr* | *nr* | *nr* | *nr* |
| CB P14, adult PC dendritic branching | N | *nt* | *nr* | - | *nr* | - |

N=normal; + = increased; - = decreased; *nr* = not reported; *nt*=not tested; DG= dentate gyrus; INP=intermediate progenitors; CA1= *cornu Ammonis* 1; HMN=hilar mossy neurons; PC= Purkinje cells; EGL=external granule layer
<sup>a</sup> Li *et al.*, 2022
<sup>b</sup> HC pathology examined in *Emx1-cre*, ex8<sup>-/-</sup>; Hori *et al.*, 2020; CB pathology in *En1-cre-cre*, ex8<sup>-/-</sup>; Yamashiro *et al.*, 2020
<sup>c</sup> Castanza *et al.*, 2021
<sup>d</sup> Weisner, Chen *et al.*, 2020; <sup>e</sup> HMN were normal in number but showed significantly reduced expression of dopamine receptor, DRD2

In contrast to CB, the mutant HC did display many of the dramatic the cellular pathologies described for C-terminal mutations. For HC, we were also able to compare *Nes-cre,* ex7^-/-^ pathologies to those found in *Emx1-cre,* ex6^-/-^ animals (Li et al., 2022). We observed significantly increased cell-packing density in the DG granule cell (gc) layer in mutant hippocampus (**Fig. 3a**), although we did not see increased packing density in *Nes-cre,* ex7^-/-^ CA1 pyramidal cell (pc) cells as reported for the 16Gso mutant (Weisner et al., 2019) (**Fig. 3a, Table 2**). We also observed decreased structural thickness in the DG gc layer in both P14 and adult animals and smaller overall hippocampal size (**Fig. 3b, Supp. Fig. 1b**), as also reported for other *Auts2* mutants including *Emx1-cre*, ex6^-/-^ mice (Li et al., 2022). Counting the number of cells stacked in layered columns in the DG, we found that the numbers of DG granule cells were indeed significantly lower compared to WT littermates in both P14 and adult animals (**Fig.3c**). In addition, the DG gc layered column structure was less well organized in P14 mutant animals than in WT, a phenotype that persisted into adulthood (data not shown). The reduced DG size thus appeared to be driven by both increased density in and a smaller total number of cells arranged in vertical columns across the DG (**Fig. 3a-c**). The increased density of the DG gc could reflect deficient dendritic growth and branching, as reported in human ASD patient postmortem studies (Amaral et al., 2008) and as also noted for *Emx1-cre*, ex8^-/-^ mice (Hori et al., 2020). On the other hand, the decreased total number of cells could reflect either a deficiency in the progenitor pool or a reduced survival of cells at later stages.

Possibly related to both the abnormal cell packing and reduced cell numbers, we documented a clear reduction in the numbers of maturing/mature CalB+ DG gc relative to immature postmitotic calretinin-positive (CalR+) cells in the mutant DG granule layer at P14 (**Fig. 3d**). This finding suggested that the transition from the CalR+ to CalB+ stage was delayed or blocked in the mutant mice. A similar finding was also documented for 16Gso mutants (Weisner et al., 2019); in contrast, P14 *Emx1-cre*, ex6^-/-^ mice were reported to display normal relative numbers of immature and mature DG cells (Li et al., 2022). Additionally, in the *Nes-cre*, ex7^-/-^ P14 HC, CalB+ neurons expressed the CalB protein at substantially reduced levels (**Fig. 3e**). As observed for *Emx1-cre,* ex6^-/-^ mutants, the main suprapyramidal axon bundle emanating from the DG gc neurons was significantly reduced in thickness in both P14 animals and adults (**Fig. 3f**) and the numbers of TBR2+ intermediate progenitors (INP) located ectopically outside the DG granule layer (**Supp. Fig. 2**) was increased at P14 suggesting abnormal migration patterns of the INP pool (**Fig. 3g; Table 2**).

*Emx1-cre*, ex15^-/-^ mice, which ablate both AUTS2 isoforms, also showed significantly reduced overall numbers of TBR2+ INP (Castanza et al., 2021), but we did not observe that phenotype (**Fig 3g**). Therefore, progenitors were normally produced and survived in the *Nes-cre,* ex7^-/-^ mice, suggesting that INP proliferation or survival, or the transition from proliferation to the post-mitotic state, are functions that require the participation of AUTS2-s. Looking beyond the DG gc layer, hilar mossy neurons (HMNs) and their calretinin-positive (CalR+) axon tracts appeared fully normal in *Nes-cre,* ex7^-/-^ HC (**Fig. 3h**), suggesting that AUTS2-s LOF played a key role in the HMN loss and disappearance of HMN axon tracts observed in *Emx1-cre*, ex15^-/-^ animals (Castanza et al., 2021).

In *Emx1-cre*, ex15^-/-^ hippocampus and in other mutants that affect both AUTS2 isoforms, hippocampal hyper-excitation, a signal of excitation/inhibition (E/I) circuit imbalance, was documented by increased expression of the immediate early gene (IEG) cFOS (Castanza et al., 2021; Hori et al., 2020; Weisner et al., 2019). However, in contrast, *Emx1-cre,* ex6^-/-^ mice showed lower numbers of c-Fos positive cells than WT controls (Li et al., 2022). Given the normal numbers and structures of HMNs, we were therefore surprised to observe a significantly increased percentage of cFOS+ cells in the *Nes-cre,* ex7^-/-^ CA1 region (**Fig. 4e**). Looking more closely at the hippocampal inhibitory circuit that includes HMNs, we found that adult mutant *Nes-cre*, ex7^-/-^ DG and CA1 regions contained significantly fewer parvalbumin-expressing (PV+) interneurons than WT littermates (**Fig. 4a**).

**Figure 4.**
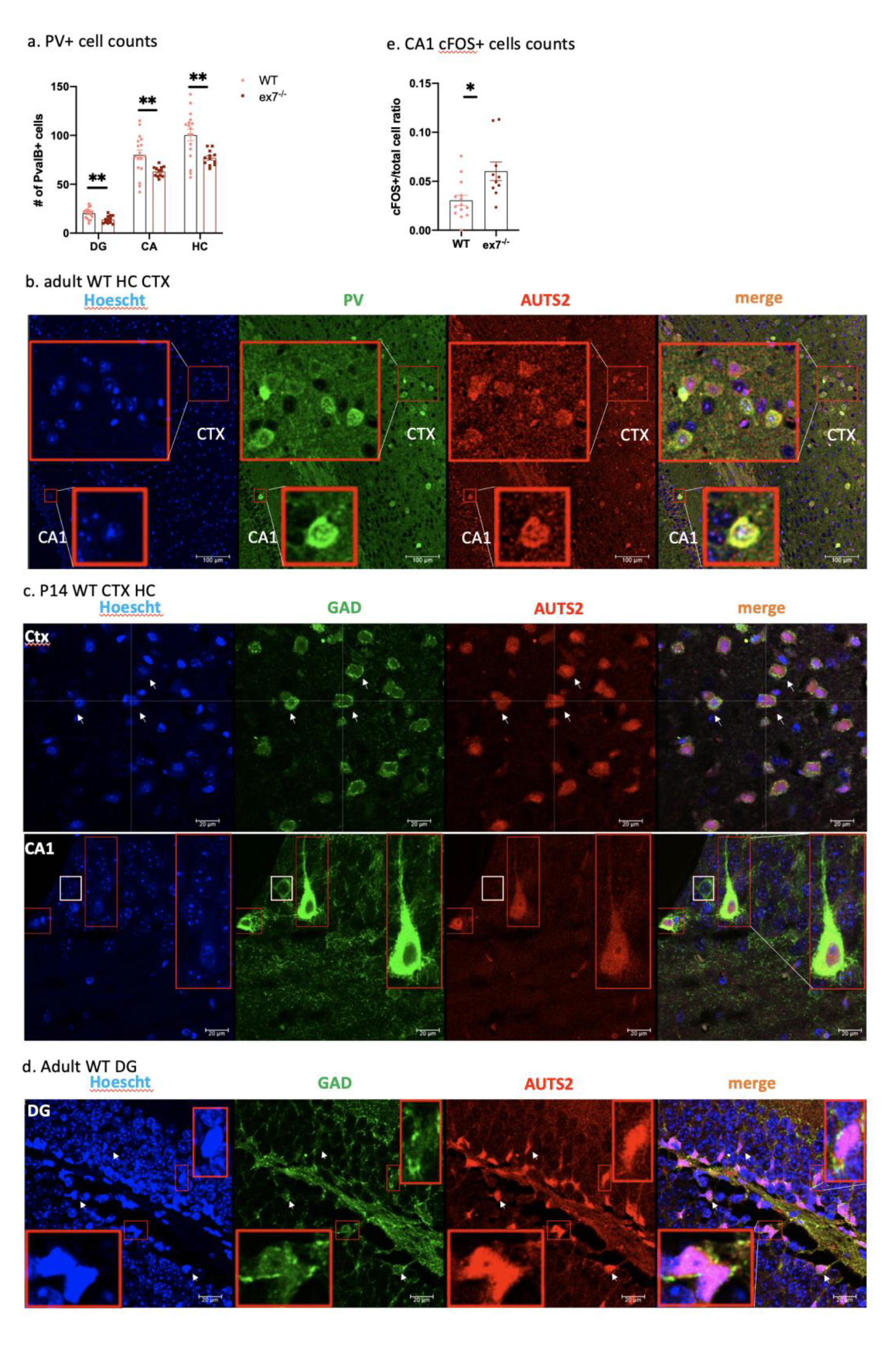
**Hippocampal PV+ interneurons express AUTS2 and are reduced in numbers in in *Nes-cre,* ex7^-/-^ mice**, abbreviated as ex7^-/-^ in figures. **a) Hippocampal PV+ cells** were significantly decreased in all HC regions in *Nes-cre,* ex7^-/-^ mutants compared to WT littermates, as were overall counts. **b)** In the cortex and HC of adult wildtype mouse, **PV+ cells (green) showed abundant AUTS2 co-expression (red)**. Red boxes show zoomed in view of indicated area. **c)** In P14 cortex and HC, some (white arrows and red zoom boxes) but not all (white box) GAD+ interneurons (green) co-expressed AUTS2 expression (red) at high levels. Cursor and white arrows show examples of overlap of GAD+ AUTS2+ cells. Red boxes show zoomed in view of indicated area. **d) In adult DG, GAD+ cells (green) showed abundant expression of AUTS2** (red). Red boxes show zoomed in view of indicated area. **e)** The ratio of cFOS+/total cell count in CA1 was significantly increased in adult mutants compared to WT. Two-tailed t-test was used to test statistical significance, * = p<0.05, ** = p<0.01, ***= p<0.001, **** = p<0.0001.

AUTS2 is most widely expressed in excitatory cells, but several studies have shown that the gene/protein is also expressed in some inhibitory interneurons, including both forebrain and cerebellum (Bedogni et al., 2010; Weisner et al., 2019; Yamashiro et al., 2020). In contrast, one report claimed that in forebrain, the protein is expressed exclusively in excitatory neurons (Hori et al., 2014). However, we found that most PV+ cells in the adult hippocampus co-expressed AUTS2 protein (**Fig. 4b**). PV first appears in cortical and hippocampal interneurons postnatally, and not all interneurons destined for PV+ fate will have made the transition by P14 (Alcántara et al., 1996). However, we also found co-expression of GABAergic marker GAD67 and AUTS2 in P14 hippocampus and cortex (**Fig. 4c**). Furthermore, although AUTS2 expression drops to low levels or disappears in most neurons as animals mature (Bedogni et al., 2010), AUTS2 protein was co-expressed with PV and GAD67 in some cells in P60 adults (**Fig. 4 b,d**). This finding is consistent with recent single nucleus sequencing data showing that *AUTS2* transcripts are most highly expressed in interneurons within the adult motor cortex (Bakken et al., 2021, summarized in the human protein atlas https://www.proteinatlas.org/ENSG00000158321-AUTS2/single+cell+type).These results suggested that AUTS2-l LOF might directly impact the development, survival, and/or mature function of inhibitory cells.

Together with published results, the data suggest that both the HMN and PV+ components of the hippocampal inhibitory circuit may depend on *Auts2* function, although the contributions of AUTS2-s and AUTS2-l in this context appear to be distinct. Further the data suggested that although HMNs survive and appear to be functional after brain-wide AUTS2-l ablation, a reduced signal from downstream PV+ interneurons could drive pathological hyper-excitation of CA1 pyramidal neurons.

### Brain-wide AUTS2-l LOF is associated with specific behavioral phenotypes

Along with brain pathologies, several dominant and recessive behavioral phenotypes have been associated with *Auts2* LOF (**Table 1**). We tested *Cmv-cre*, ex7^-/+^ animals to identify dominant phenotypes and the more robust *Nes-cre,* ex7^-/-^ adults to test recessive phenotypes. We focused on behavioral tests that have also been used to study key behaviors in other *Auts2* mutant mice. For example, we found novel object recognition (NOR) deficits, in the *Nes-cre,* ex7^-/-^ mice, but only as a recessive trait (**Fig. 5d; Table 1; Fig. 6a**); NOR deficits are dominantly expressed in *Auts2^neo/+^* (Hori et al., 2015) and *Auts2^del8/+^*(or *Cag-cre,* ex8^-/+^), which carry a germline deletion of *Auts2* exon 8 (Hori et al., 2014, 2020). Here we should also note a contrast with a recent study reporting that *Emx1-cre,* ex6^-/-^ mice, (which also ablate AUTS2-l in forebrain excitatory neurons) *did not* display NOR deficits (Li et al., 2022). In further contrast to *Emx1-cre,* ex6^-/-^ mice, we found reduced sociability in *Nes-cre,* ex7^-/-^ animals (**Fig. 5e; Table 1**) but not deficits in social memory (recognizing a familiar mouse) (**Fig. 5f; Table 1**). We cannot presently explain these conflicting results but hypothesize that the contrasts may reflect the activities of the different *cre* alleles.

**Figure 5.**
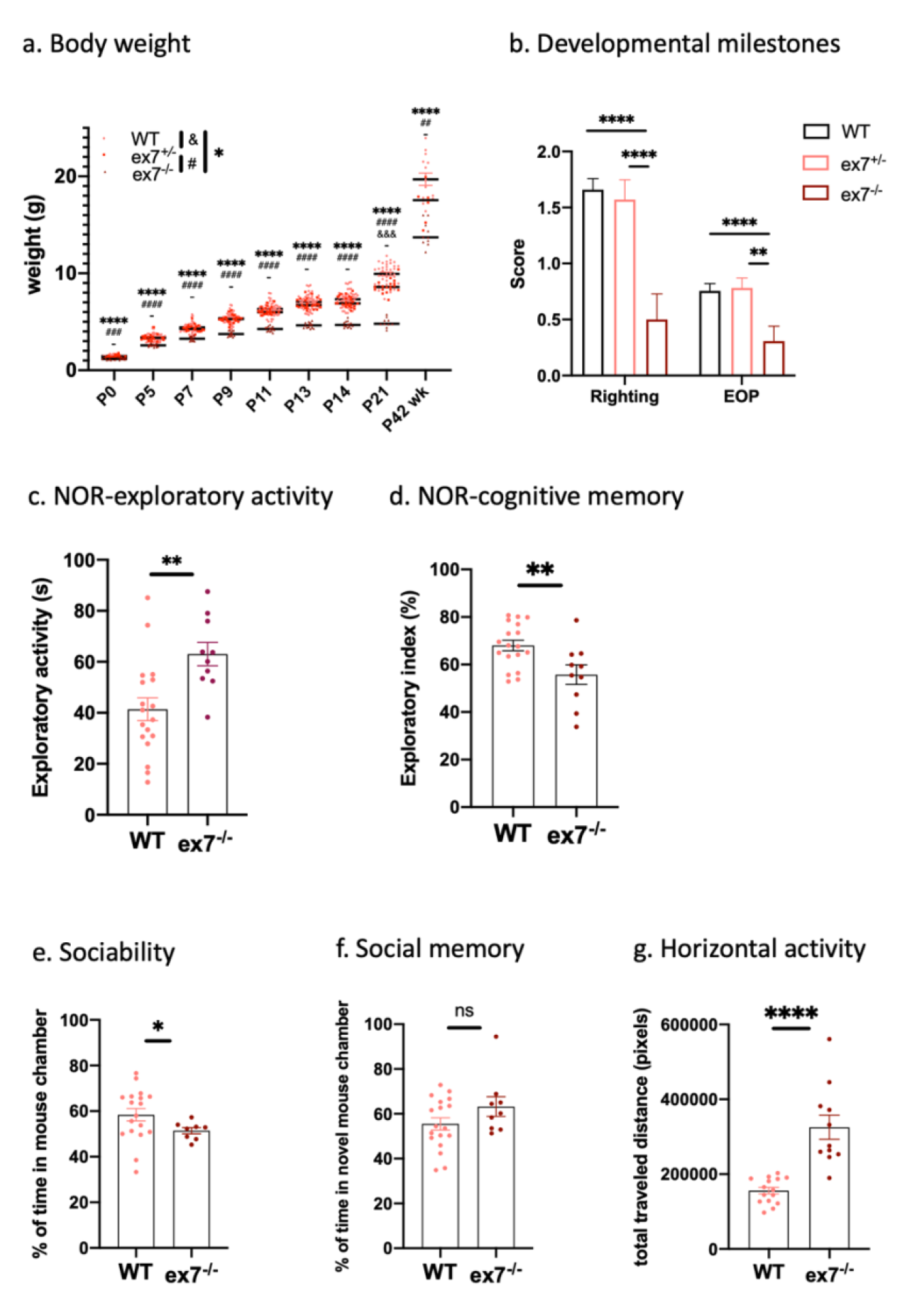
**Developmental and behavioral characterization of *Nes-cre*, ex7^-/-^ mutants**, abbreviated as ex7^-/-^ in figures. **a) Body weight measurements at different developmental time points.** P0, P5, P7, P9, P11, P13, P14, and P21 pups and adults (P42-P49) were measured for body weight. For P0 measurements, *Cmv-cre*, ex7^-/-^ and *Cmv-cre*, ex7^-/+^ were used, and all other measurements were with *Nes-cre*, ex7^-/-^ and *Nes-cre*, ex7^-/+^. The ex7^-/-^ mice weighed significantly less than both WT and ex7^-/+^ mice on all measured time points, and ex7^-/+^ mice weighed significantly less than WT mice on P21. WT n= 15-49; ex7^-/+^ n = 6-25; ex7^-/-^ n= 10-17. **b) developmental delay tests of righting reflex and eye-opening (EOP)** scores. Scores of 0-2 were assigned (see Methods), and *Nes-cre*, ex7^-/-^ mice showed delay in both righting reflex and EOP test compared to WT mice. WT n=44-45; *Nes-cre*, ex7^-/+^ n = 21-24; *Nes-cre*, ex7^-/-^ n= 14. **c,d) Total exploratory activity during NOR test training session and exploratory index during retention session**. The *Nes-cre*, ex7^-/-^ mice showed significant increase exploratory activity in training= session and decreased exploratory index in retention session. WT n= 18; *Nes-cre*, ex7^-/-^ n= 11. **e,f) Sociability and Social Memory**. **The Sociability** panel shows the percentage of time a tested mouse spent in mouse chamber vs. total time spent in mouse chamber plus the toy chamber during second 10 min session of three-chamber test (see method). **The social memory** panel shows the percentage of time a tested mouse spent in novel mouse (stranger mouse 2) chamber vs. total time spent in familiar mouse (stranger mouse 1) chamber plus novel mouse (stranger mouse 2) chamber. The *Nes-cre*, ex7^-/-^ show significantly decreased sociability but normal social memory compared to WT mice. Sociability: WT n=18; *Nes-cre*, ex7^-/-^ n=9, within which 1 outlier excluded (through Grubbs’test G=2.543 and with ROUT test Q=1%); Social memory: WT n= 18; *Nes-cre*, ex7^-/-^ n=9, within which 1 data point can be excluded through Grubb’s test as outlier G= 2.372, but same value cannot be excluded with ROUT test; excluding the data point does not change result significance. **g) total horizontal activity** in unit of pixels after normalization. Total distance traveled horizontally was recorded and quantified as described in method. *Nes-cre*, ex7^-/-^ show significantly increased horizontal activity compared to WT mice. WT n= 15; *Nes-cre*, ex7^-/-^ =11. Two-tailed t-test was used to test statistical significance, * = p<0.05, ** = p<0.01, *** = p<0.001, **** = p<0.0001, ns = not significant.

**Figure 6.**
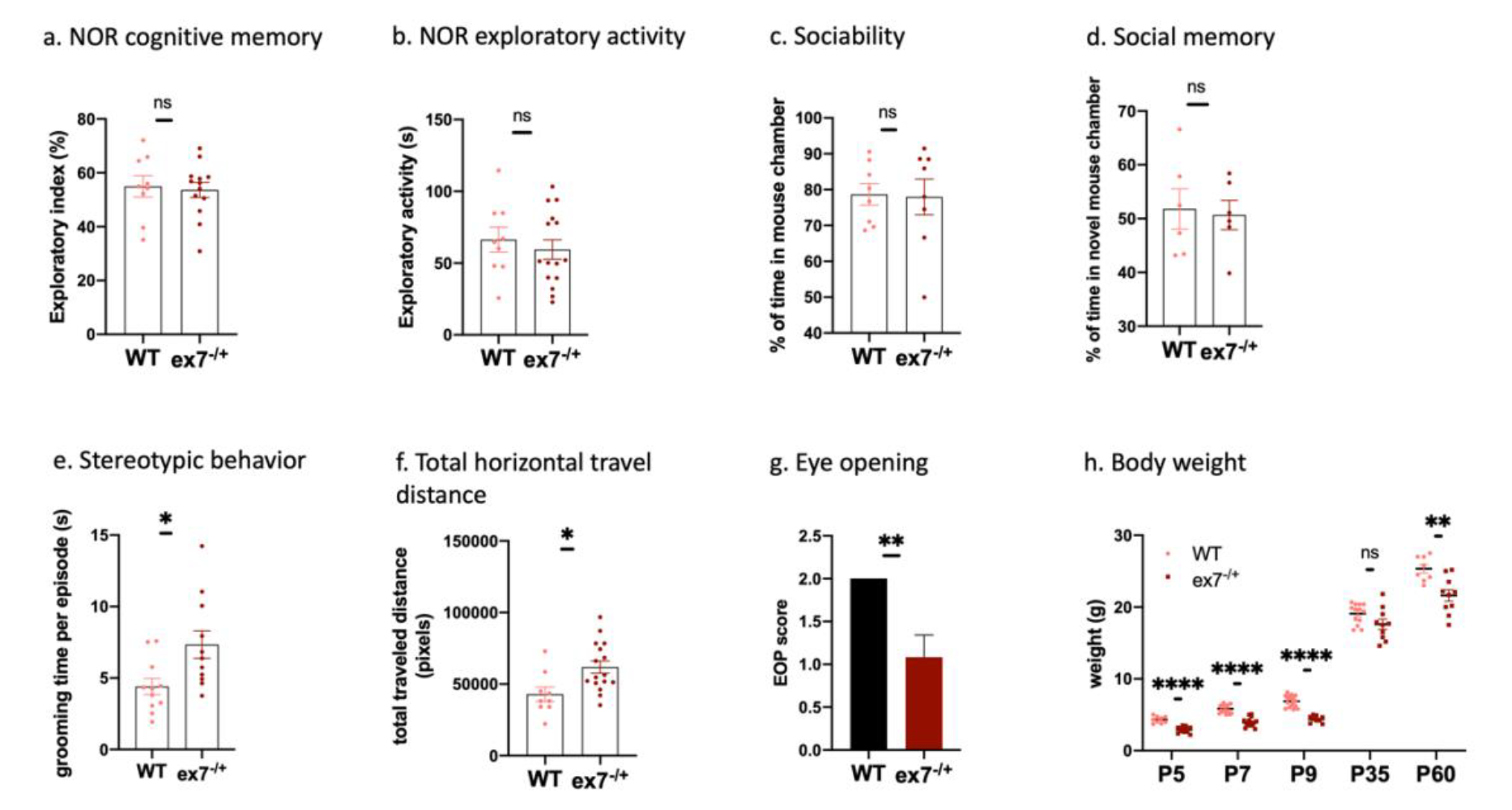
**Behavioral characterization of *Cmv-cre*, ex7^-/+^ animals,** abbreviated as ex7^-/+^ in figures. **a) NOR-cognitive memory in *Cmv-cre*, ex7^-/+^** animals compared to WT controls show no significant difference. WT n=9, *Cmv-cre*, ex7^-/+^ n = 13; **b) NOR-exploratory activity in *Cmv-cre*, ex7^-/+^** animals compared to WT controls show not significant difference. WT n=9, *Cmv-cre*, ex7^-/+^ n = 16. 1 outlier removed from mutant group (through Grubb’s test, G=2.903, Alpha = 0.05). **c) Sociability in *Cmv-cre*, ex7^-/+^** animals compared to WT controls show no significant difference. WT n=8, *Cmv-cre*, ex7^-/+^ n = 8. **d) Social memory in *Cmv-cre*, ex7^-/+^**animals compared to WT controls show no significant difference. WT n=6, *Cmv-cre*, ex7^-/+^ n = 6. **e) Stereotypic behavior in *Cmv-cre*, ex7^-/+^** animals show significant increase compared to WT animals. WT n=11, *Cmv-cre*, ex7^-/+^ n = 11. **f) Total horizontal travel distance of *Cmv-cre*, ex7^-/+^** animals compared to WT controls show significant increase. WT n=9, *Cmv-cre*, ex7^-/+^ n = 16.**g) Developmental milestone – eye opening was delayed in *Cmv-cre*, ex7 ^-/+^ animals** compared to WT controls. WT n=13, *Cmv-cre*, ex7 ^-/+^ n=12. **h) Body weight measurements on *Cmv-cre*, ex7 ^-/+^ animals** compared to WT controls show mutant animals had significantly decreased body weight at different developmental time points. WT n=8-17, *Cmv-cre*, ex7 ^-/+^ n=9-15. Two-tailed t-test was used to determine statistical significance; ns = p>0.05, * = p<0.05, ** = p<0.01, **** = p<0.0001, ns = not significant.

We found that *Cmv-cre*, ex7^-/+^ but not *Nes-cre,* ex7^-/+^ mice showed dominant delay in achieving developmental milestones (**Fig. 6g, 5b**). Other dominant phenotypes associated with ex7 deletion included reduced postnatal growth (both *Cmv-cre* and *Nes-cre* heterozygotes) (**Fig. 5a, 6h**), and excessive stereotypic grooming behaviors (in *Cmv-cre,* ex7^-/+^ mice) (**Fig. 6e**). Excessive grooming behaviors were also observed for *Emx1-cre,* ex6^-/-^ mice (Li et al., 2022). Noting a phenotype that has not been associated with *Auts2* cKO of any type in the past, we quantified total traveling distance in the open field test and confirmed that both heterozygous *Cmv-cre,* ex7^-/+^ and homozygous *Nes-cre, ex7^-/-^* animals were significantly more active than WT littermates (**Fig. 5g**, **Table 1, Fig.6f**). Hyperactivity was also reported for 16Gso heterozygotes, although homozygotes of that genotype were less active than WT littermates(Weisner et al., 2019). Furthermore, this hyperactive phenotype stood in stark contrast to the reduced activity levels reported for *Auts2^neo/+^* and *Cag-cre*, ex8^-/+^ mice (Hori et al., 2015, 2020). The results suggested that AUTS2-l LOF contributes centrally to a hyperactive phenotype, but also suggested that AUTS2-s might modulate the expression of this trait.

### **Targeting AUTS2-l LOF to the *Calb1*-lineage yields specific subsets of *AUTS2-*linked phenotypes**

Published data suggest a role for AUTS2-l in neuron migration and maturation, functions that could possibly be executed in both excitatory and inhibitory neurons. Since mature DG gc express CalB and several types of hippocampal interneurons, including PV+ cells, express CalB a least transiently during their development (Alcántara et al., 1996), we could target both cell populations with the *Calb1-cre* allele. We tested the effects of AUTS2-l LOF in these cell types by generating *Calb1-cre, ex7^-/-^* animals and examining morphology, behavior, and brain pathology in comparison to WT littermates.

Young *Calb1-cre,* ex7^-/-^ mice showed delayed establishment of the righting reflex (**Fig. 7a**), but they were not smaller than WT littermates **(Suppl. Fig. 3a)** in contrast to animals with germline or brain-wide ex7 cKO, or to animals in which AUTS2-l was ablated more broadly in forebrain excitatory neurons or in mature excitatory cells (*Emx1-cre* or *Camk2a-cre*, ex6^-/-^ genotypes; (Li et al., 2022)). Furthermore, *Calb1-cre, ex7^-/-^* mice showed no difference from WT littermates in terms of social preference **(Suppl. Fig 3b),** or levels of exploratory activity (**Fig. 7b**). On the other hand, they displayed clear deficits in NOR (**Fig. 7c**) and hyperactivity in the open field test compared to WT littermates (**Fig. 7d**). Therefore, the ablation of AUTS2-l in CalB+ cells alone was sufficient to recapitulate the cognitive deficits and hyperactivity seen in brain-wide AUTS2-l KO animals, but not reduced social interaction or other key mutant phenotypes.

**Figure 7.**
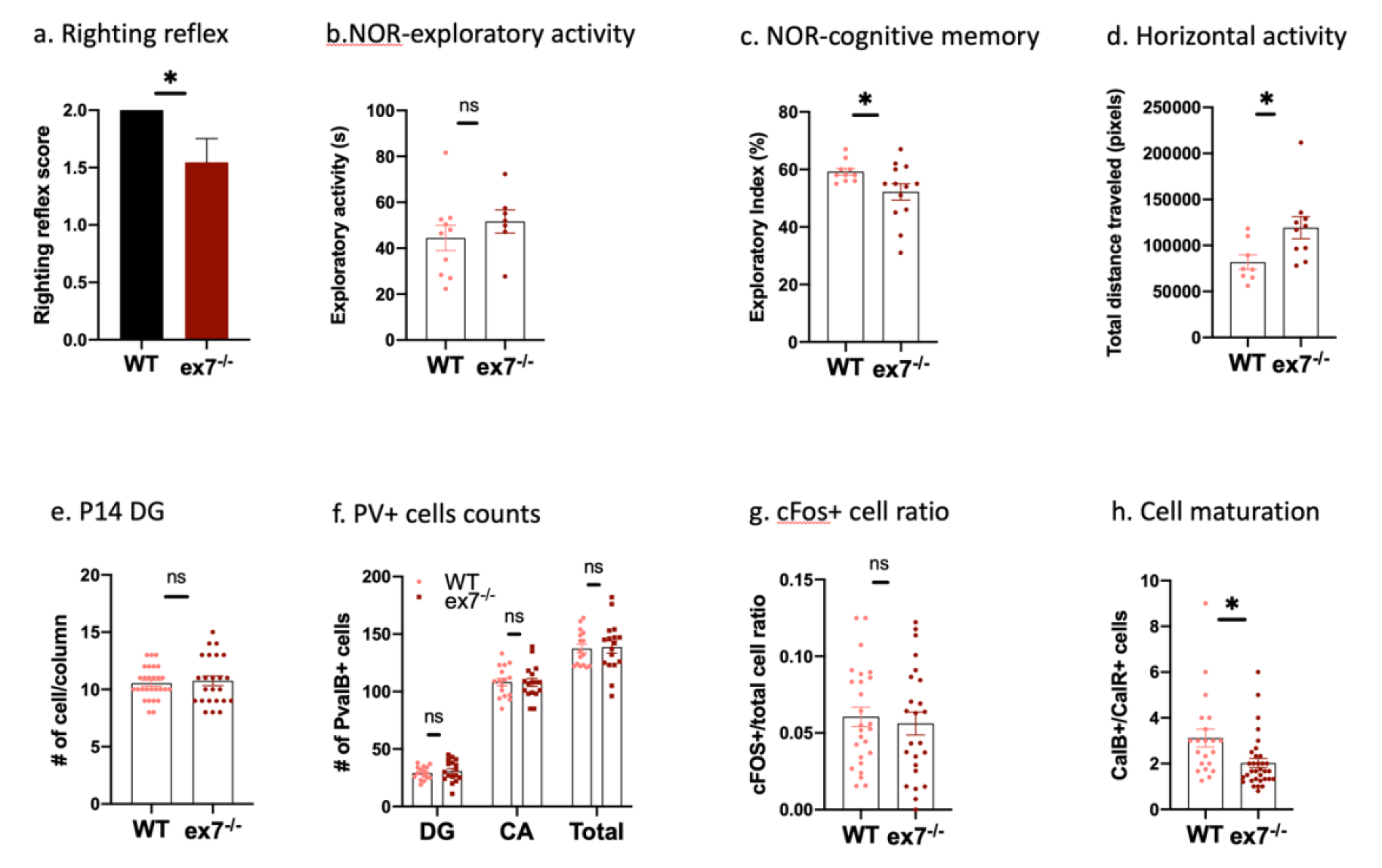
**Developmental and behavioral characterization of *Calb1-cre*, ex7^-/-^ mutant**, abbreviated as ex7**^-/-^** in figures. **a) Righting reflex scores**. Scores of 0-2 were assigned according to method and ex7**^-/-^** mice showed delay in righting reflex compared to WT mice. WT n=13; *Calb1-cre*, ex7^-/-^ n= 11; **b, c) Total exploratory activity** during NOR test learning session and exploratory index during the test session. The *Calb1-cre*, ex7^-/-^ mice did not show significant difference in total exploratory activity in training session compared to WT mice but showed significantly decreased exploratory index in retention session. WT n= 10-11; *Calb1-cre*, ex7^-/-^ n= 7-13. 1 outlier removed from WT group (through Grubbs’ test, G= 2.434, alpha = 0.05 and with ROUT test Q=1%). **d) Total horizontal distance traveled in open field test.** *Calb1-cre*, ex7^-/-^ mice showed significantly increased in total horizontal travel distance compared to WT mice. WT n=8; *Calb1-cre*, ex7^-/-^ n= 10. **e) number of cells stacking per column in P14 DG** of mutant compared to WT animals showed no significant difference. **f**) PVALB+ cells in different regions of HC were not significantly different in mutant compared to wildtype animals as well as sum counts. **g)** The ratio of cFOS+/total cell count in CA1 was not significantly different in adult mutants compared to WT. **h) The ratio of CalB+/CalR+ cells** was significantly decreased in ex7^-/-^ mutants HC DG compared to WT. Two-tailed t-test was used to test statistical significance, * = p<0.05, ** = p<0.01, *** = p<0.001, **** = p<0.0001, ns = not significant.

Looking at brain pathology, we found that in contrast to other *Auts2* mutants including C-terminal mutants, *Nes-cre,* ex7^-/-^ and *Emx1-cre,* ex6^-/-^ animals, the hippocampus of *Calb1-cre*, ex7^-/-^ mice was not smaller than that of normal littermates, and the total numbers of cells in the DG granule layer was not reduced (**Fig. 7e**). This finding may be related to the fact that TBR2+ cells in *Calb1-cre*, ex7^-/-^ hippocampus were distributed normally and were not ectopically positioned (**Suppl. Fig3c**), as they were in *Nes-cre,* ex7^-/-^ mice. Furthermore, we found that PV+ cell counts were *not* decreased in the *Calb1-cre* mutants compared to littermate controls (**Fig. 7f**). To ask whether there might still be a malfunction in hippocampal inhibitory circuits signaled by hyperexcitation, we tested c-Fos expression in *Calb-cre,* ex7^-/-^ mice but found no difference between the mutants and WT littermate controls (**Fig.7g**).

Therefore PV+ neuron loss is correlated with the c-FOS associated hyper-excitation phenotype, but the *Calb1-cre,* ex7^-/-^ animals displayed NOR deficits without either of these E/I phenotypes. Furthermore, the data suggest that the loss of PV+ IN was not driven by AUTS2-l LOF in the interneurons themselves, at least not after the cells express *Calb1-cre* in late gestation. On the other hand, as in *Nes-cre*, ex7^-/-^ animals, the ratio of CalB/CalR+ neurons in the DG granule layer was significantly lower in the *Calb1-cre*, ex7^-/-^ mutants relative to WT littermates (**Fig. 7h**). This finding indicated that the delay or failure of DG gc to fully mature was driven by AUTS2-l LOF within the DG gc themselves, and that this block was centrally related to expression of NOR deficits.

### Disturbed expression of neurodevelopmental genes after AUTS2-l LOF

AUTS2 has been implicated in chromatin remodeling, regulation of transcription and RNA binding/stability, all of which can be predicted to impact RNA transcription. Indeed, *Auts2* chromatin-binding assays (Gao et al., 2014; Oksenberg et al., 2014) and transcriptomic analyses involving *Auts2* mutants of various types (Castanza et al., 2021; Hori et al., 2020; Li et al., 2022; Liu et al., 2021; Weisner et al., 2019) have confirmed a role in regulating the expression of neurodevelopmental genes. However, transcriptomic changes have not been examined after brain-wide AUTS2-l LOF *in vivo.* With the goal of linking transcriptomic changes due to AUTS2-l LOF to behavioral and brain pathologies, we carried out RNA-seq comparing gene expression in mutant P0 FB (*Cmv-cre*, ex7^-/-^), and P14 HC (*Nes-cre*, ex7^-/-^) to samples from WT littermates of both ages (**Fig. 8**). Hundreds of differentially expressed genes (DEGs) were identified in both sample sets, although P0 and P14 DEGs showed relatively little overlap: only 56 genes were commonly detected among the 397 P0 and 546 P14 DEGs. Furthermore, more than half of these overlapping DEGs (31 genes) were oppositely expressed at P0 and P14 (**Supp. Table 1**). DEGs at both time points were strongly enriched in shared functional categories/pathways related to neuron differentiation and function, but like the shared DEGs, several shared pathways were oppositely up-or down-regulated at the two stages (**Table 3**; **Fig. 8**).

**Figure 8.**
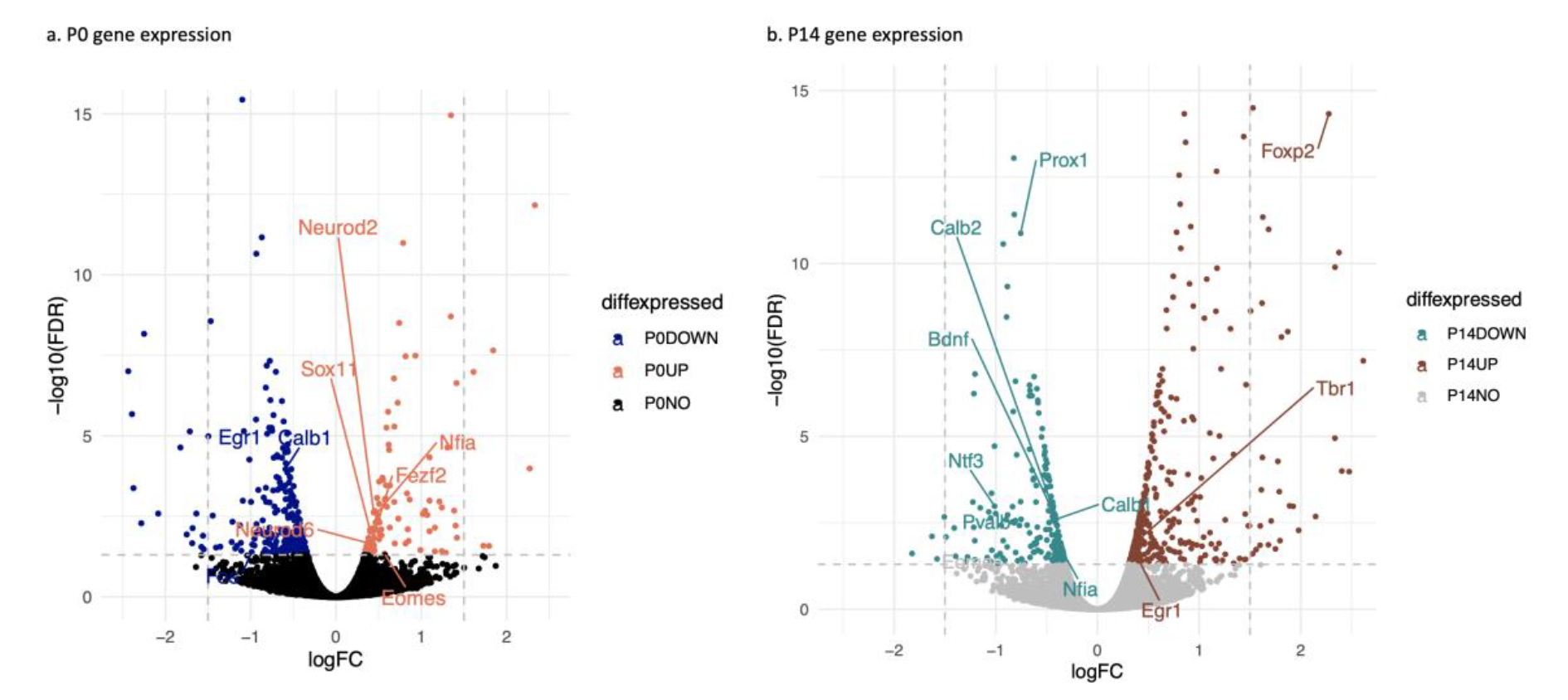
**Gene expression volcano plot of RNA-seq data from P0 FB and P14 HC ex7^-/-^ compared to WT controls**. **a) P0 gene expression**, blue = down-regulated genes, coral = up-regulated genes, black = not differentially expressed; **b) P14 gene expression**, green = down-regulated genes, brown = up-regulated genes, grey = not differentially expressed. For both P0 and P14 plots, view is zoomed to regions with –log10 (FDR) between 0-15, and log FC between −2.5 to 2.5. Grey dashed lines represents log FC = −1.5, 1.5, and FDR = 0.05; Plots with all genes are in Supplementary figure 3 and gene expression data are fully presented in Supplementary Table 1.

**Table 3.** Functional category enrichments for DEGs from *Auts2-ex7^-/-^* P0 forebrain and P14 hippocampus.

| **Functional category** | **Enrichment -logP** |  |  |  |
| --- | --- | --- | --- | --- |
|  | **P0 FB** |  | **P14HC** |  |
|  | **<u>up</u>** | **<u>down</u>** | **<u>up</u>** | **<u>down</u>** |
| GOMF DNA-binding transcription activator activity, RNA polymerase II-specific | 10 | - | - | - |
| GOMF ion channel activity | - | 10 | 5.2 | - |
| GOMF glutamate receptor activity | - | 10 | - | - |
| GOMF adenylate cyclase inhibiting G protein-coupled glutamate receptor activity | - | 5.63 | - | - |
| GOMF extracellular matrix structural constituent | - | - | - | 10 |
| GOMF nerve growth factor receptor binding | - | - | - | 4.86 |
| GOBP positive regulation of transcription by RNA polymerase II | 10 | - | - | - |
| GOBP forebrain development | 5.1 | - | 10 | 5.08 |
| GOBP axonogenesis | - | 5.71 | 10 | 5.57 |
| GOBP cell morphogenesis involved in neuron differentiation | - | 10 | 10 | 10 |
| GOBP trans-synaptic signaling | - | 10 | 10 | - |
| GOBP learning or memory | - | 10 | 10 | 5.32 |
| GOBP regulation of membrane potential | - | 10 | 10 | - |
| GOBP neurotransmitter transport | - | - | 10 | - |
| GOBP neurotransmitter secretion | - | 10 | 10 | - |
| GOBP glial cell differentiation | - | - | - | 5.89 |
| GOBP blood vessel development | - | - | - | 5.6 |
| GOBP synaptic transmission, glutamatergic | - | 5.75 | 10 | - |
| GOBP synaptic vesicle exocytosis | - | 5.92 | 5.73 | - |
| GOBP negative regulation of neurogenesis | - | - | - | 5.13 |
| GOBP Pallium development | - | - | 5 | - |
| GOBP axon guidance | - | - | 10 | 5.93 |
| GOBP cellular response to growth factor stimulus | - | - | 5.81 | 5.09 |
| GOBP synapse organization | - | - | 10 | - |
| GOBP dendrite development | - | - | 10 | - |
| GOBP behavioral fear response | - | - | 10# | - |
| GOBP regulation of cell migration | - | - | - | 10 |
| GOBP regulation of neuron differentiation | - | - | - | 10 |
| GOBP negative regulation of neurogenesis | - | - | - | 5.13 |
| MGI abnormal cognition | - | 10 | 10 | - |
| MGI abnormal synaptic physiology | - | 10 | 5.41 | - |
| MGI abnormal learning/memory/conditioning | - | 10 | 10 | - |
| MGI abnormal brain interneuron morphology | - | 5.13 | - | - |
| MGI abnormal dentate gyrus morphology | - | - | - | 10 |
| MGI abnormal neuron morphology | - | - | - | 10 |

Functional enrichments were determined for up- and down-regulated $\text{fdr} < 0.05$ DEGs from comparisons of *Aut2-ex7* mutant P0 FB (forebrain) and P14 HC (hippocampus) to WT littermates as described in the text, using the ToppCluster Suite (cite). Categories shown were selected to be highly enriched, but more specific categories to represent groups of related/overlapping functional categories. In the ToppCluster output, all categories with $-\log P \geq 10$ are listed with a value of 10 (in columns 3-6). GOBP= GO biological process, GOMF= GO molecular function, MGI= mouse genome informatics Mouse Phenotypes.

Interested in understanding how this functional shift might be regulated, we focused on differentially expressed transcription factors (TF) and chromatin regulators and found genes with well-established roles in forebrain and hippocampal development. For example, among the P0 up-/P14 down-regulated group was *Sox11,* which promotes the proliferation of the excitatory neuron precursors in developing cortex and hippocampus and inhibits their differentiation (Hoshiba et al., 2016; Liu et al., 2019; Wang et al, 2013) and later promotes hippocampal DG gc polarity and dendrite formation (Abulaiti et al., 2022; Hoshiba et al., 2016). Concordant with the pattern of *Sox11* expression, genes associated with neuron differentiation and mature neuron function including neurite and synapse formation were *down-regulated* in P0 FB. However, “DNA-binding transcriptional activator activity” was the single most highly enriched functional category for P0 up-regulated DEGs (**Table 3**) and included well known regulators such as *Eomes* (encoding TBR2)*, Insm1, Neurod2, Neurod6,* and Notch-related TF, *Rbpj* (**Fig. 8**).

In contrast to P0, genes involved in mature-neuron functions such as neurotransmitter transport and secretion were robustly *up-regulated* in the P14 hippocampus although functions related to neurite development were enriched for both up-and down-regulated DEGs. A closer look revealed that the up-regulated genes in this category specialize in the development of synapses and dendritic spines (*Foxo6, L3mbtl1, Mef2c, Nfia, Nfib, Pcdh10, Tbr1, Nrxn2, Stx1a, Cit, Cobl, Met*) while the down-regulated genes are involved more generally in DG gc differentiation and maturation (*Lef1, Neurod1*, *Neurog2, Prox1, Sox11*), in neurite differentiation and outgrowth (including growth factors, *Bdnf, Ntf3,* and *Ngf)* in axon pathfinding (*Sema3c, Sema5a, and Slit2*) and in cell migration (*Cxcl12, Lef1, Prox1*) (**Table 3; Supp. Table 1**). The *Pvalb* gene was also significantly down-regulated, consistent with the finding of reduced PV+ cell numbers in mutant P14 HC. The data were consistent with published pathology for C-terminal mutants, including the repressed growth of axons and dendrites, but excessive dendritic spine formation (Hori et al., 2020). The data were also consistent with the observation of reduced DG gc numbers but increased c-Fos levels in other published mutants as well as the *Nes-cre,* ex7^-/-^ hippocampus (**Fig. 4e**). The transcriptomic data also pointed to some possible molecular drivers of these pathological phenotypes.

### Association of P0 and P14 DEGs with AUTS2 chromatin binding peaks reveals a high correlation and potential direct target genes

To understand how AUTS2-1 LOF might impact these gene expression patterns, we looked for overlaps between DEGs and AUTS2 DNA binding peaks. Two chromatin immunoprecipitation (ChIP) datasets are publicly available for AUTS2, both generated with antibodies detecting AUTS2- and AUTS2-s isoforms together in chromatin from embryonic day 16.5 forebrain (E16.5 FB) (Oksenberg et al., 2014) or P4 whole brain (Gao et al., 2014), respectively. We reanalyzed both datasets and found a significant overlap (775 directly overlapping peaks of approximately 5000 peaks total for each set, log *p=*1E-1611, hypergeometric test). However, most peaks in each dataset mapped to unique positions, suggesting that AUTS2 DNA binding patterns might depend on developmental stage and/or brain region. Examining associations between DEGs from P0 FB and P14 hippocampus of *Nes-cre,* ex7^-/-^ mice and peaks from each dataset, we found a significant enrichment of ChIP peaks of both types located in regions that were either directly flanking or within DEGs at P0 and P14 (**Table 4**). The highest enrichment between peaks and DEGs at P0 was with *down*-regulated DEGs, consistent with the idea that AUTS2-l serves as a component of an activating complex. On the other hand, in P14 hippocampus, up-regulated DEGs were more likely to be associated with AUTS2 peaks (**Table 4**). In these and other respects, the E16.5 and P4 AUTS2 ChIP peak sets gave qualitatively similar results; because of the stronger association with DEGs, we will focus this discussion on overlaps with the E16.5 data set.

**Table 4.** Hypergeometric *p* values for overlaps between published AUTS2 ChIP peaks and DEGs.

| DEG set | E16.5<br>FB<sup>a</sup> | # DEGs | P4<br>whole<br>brain<sup>b</sup> | # DEGs | E16.5/<br>P4<br>overlap | #DEGs |
| --- | --- | --- | --- | --- | --- | --- |
| P0 FB -all | 2.22E-27 | 100 | 5.66E-27 | 138 | 1.64E-05 | 26 |
| P0 FB -down | 9.92E-25 | 78 | 7.17E-20 | 98 | 1.09E-02 | 14 |
| P0 FB -up | 2.12E-05 | 22 | 3.66E-09 | 40 | 1.79E-05 | 12 |
| P14 HC -all | 5.97E-49 | 156 | 7.64E-38 | 193 | 2.90E-02 | 22 |
| P14 HC -down | 1.11E-14 | 51 | 2.47E-09 | 60 | 5.71E-02 | 9 |
| P14 HC -up | 8.08E-37 | 105 | 1.14E-31 | 133 | 8.79E-02 | 13 |

<sup>a</sup> From Oksenberg *et al.*, 2014; <sup>b</sup> From Gao *et al.*, 2014

Two issues complicate the interpretation of these results in terms of AUTS2-l regulatory functions. First, as we have shown, AUTS2-s is up-regulated in ex7^-/-^ mutants in P0 FB but not in P14 HC (**Fig. 1**) and both isoforms could participate in chromatin binding. Second, the collection of peaks nearest DEGs included promoter-proximal and more distally located intronic and intergenic peaks; most DEG-associated peaks were distally located. Examples included peaks within or flanking neurogenesis regulators *Nfia* and *Sox11* that overlapped with ENCODE-validated long-range enhancers for those genes (Gorkin et al., 2020) (**Table 5**). Notably in this regard, *Nfia* and *Sox11* were among the 31 DEGs detected in both P0 and P14 datasets but oppositely regulated at the two time points (see above).

**Table 5.**
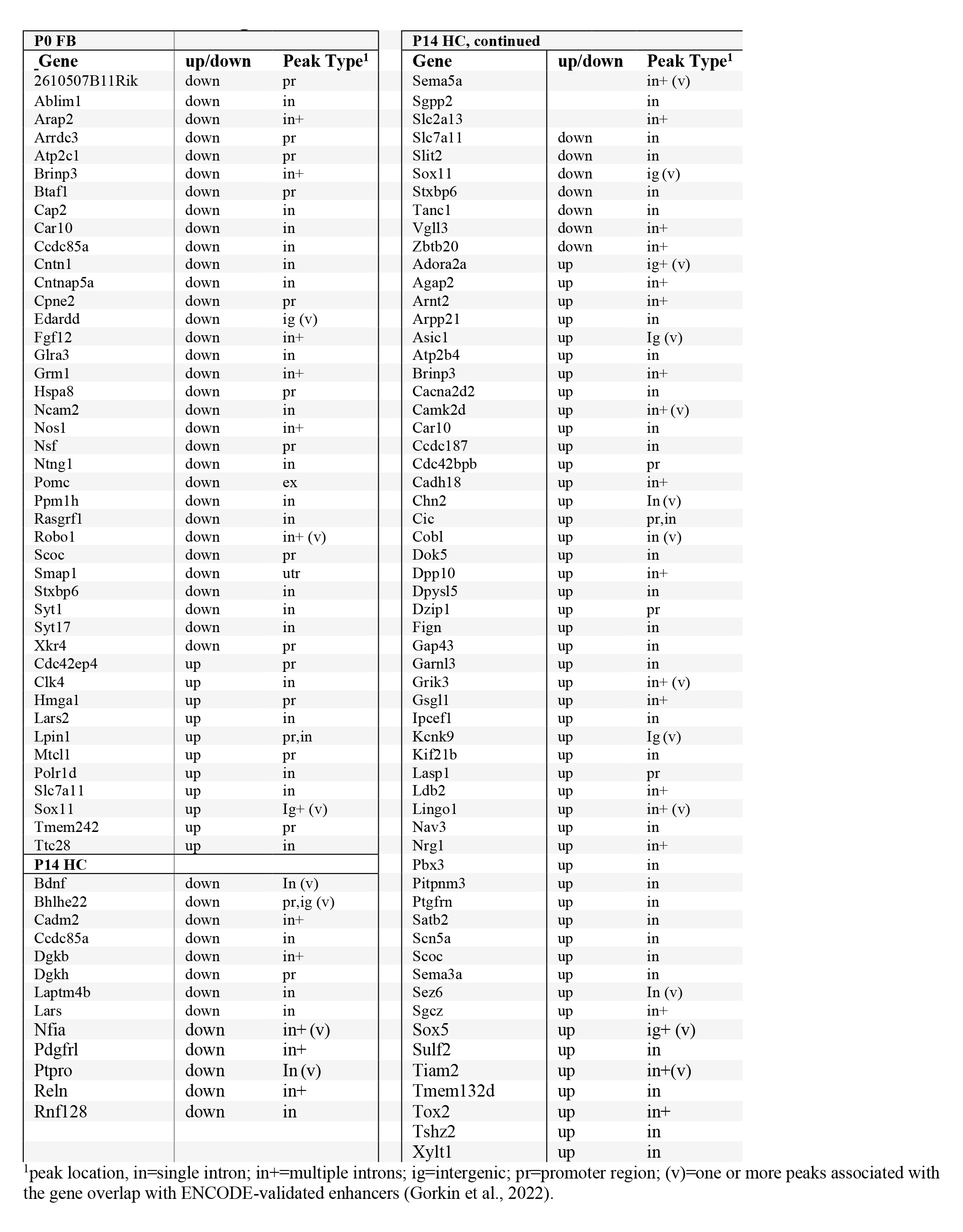
DEGs associated with E16.5 forebrain AUTS2 peaks located within the gene or intergenic but validated as DEG gene-linked enhancers.

In addition to these genes, other DEGs at both time points contained distal AUTS2 peaks within introns or in intergenic locations including validated DEG-linked enhancers (**Suppl Table 2; Table 5**). For example, E16.5 AUTS2 peaks included a peak within the neighboring *Alpk1* gene that is an ENCODE-validated distal enhancer for P14 down-regulated DEG *Neurog2,* two intergenic enhancers validated for up-regulated DEG *Sox5*, and intronic enhancers within both up-and down regulated DEGs (Table 5). Of interest, a number of ChIP peak-associated DEGs are established ASD candidate genes (*Cic, Mef2c, Meis2*, *Rims1, Reln, Rfx3, Sox5*, *Tbr1, Tle3, Tshz3, Zfp462, Zbtb20*; https://gene.sfari.org/). Other putative target genes including *Sox11* (Tsurusaki et al., 2014) are associated rare disorders that include ASD, ID, developmental delay, language disorders and other phenotypes that overlap with AUTS2 syndrome. The data thus suggest a direct role for AUTS2-l in regulating genes implicated in ASD risk.

## Discussion

These data associate the isolated loss of AUTS2-l with specific subsets of *Auts2-*linked behavioral, pathological, and transcriptomic phenotypes for the first time, revealing new information about the *in vivo* functions of the long isoform, and several new hints regarding short AUTS2 isoform functions as well. Specifically, we found deficits in both cognitive memory and social interaction in animals homozygous for brain-wide deletion of *Auts2* exon 7 but not in heterozygotes; these same deficits are expressed as dominant traits in animals carrying C-terminal mutations, indicating an important role for AUTS2-s in the expression of cognitive and social phenotypes. These findings are consistent with the relatively mild expression of AUTS2-syndrome phenotypes in human patients carrying N-terminal variants including mutations within exons 6 and 7 (Beunders et al., 2015, 2013; Sengun et al., 2016), compared to the severe ID and ASD phenotypes associated with heterozygosity for C-terminal alleles (Beunders et al., 2013, 2016; Liu et al., 2021). Also consistent with human patients, exon 7 deletion and C-terminal mutant heterozygotes display some common dominant phenotypes, including growth retardation and developmental delay, indicating a central role for AUTS2-l in the expression of those traits. Additionally, we report one striking contrast between N-and C-terminal *Auts2* LOF alleles: exon 7 deletion - whether generated with germline or brain-wide *cre* alleles – is associated with a marked, dominant hyperactivity, whereas C-terminal heterozygotes have been reported to display reduced activity levels compared to wild type mice (Hori et al., 2015).

ADHD has been proposed as a central AUTS2 syndrome phenotype (Biel et al., 2022; Sanchez-Jimeno et al., 2021b), and the mechanisms that drive this distinction between N-terminal and C-terminal *Auts2* alleles are thus of special interest. Although additional work will be required to solve this puzzle, we hypothesize that this striking difference might relate to the transient over-expression of AUTS2-s in the *ex7^-/-^*mutant neonatal brain. In this scenario, over-expression of AUTS2-s during late gestation or early postnatal life could drive the development of hyperactivity, with the reduced expression of AUTS2- s in C-terminal mutants leading to the opposite phenotype. Considering how this pathological over-expression of AUTS2-s might be actuated, we note that the group reporting AUTS2 ChIP in E16.5 forebrain chromatin found multiple AUTS2 chromatin binding sites within the *Auts2* gene (Oksenberg et al., 2014); our remapped data in the mm10 genome build confirmed significant peaks in introns 2,3,4, 5 and 13 (**Supp. Table 2**). These internal AUTS2 binding peaks were not detected in P4 whole brain ChIP (Gao et al., 2014), suggesting the possibility of a transient suppression of AUTS2-s encoding transcripts by AUTS2-l binding. Published data have shown that the AUTS2 isoforms act sequentially during neurogenesis, with AUTS2-l arising after the initial expression of AUTS2-s in neuron progenitors both *in vitro* (Monderer-Rothkoff et al., 2021) and *in vivo* (Castanza et al., 2021; Liu et al., 2021) to instigate neuron maturation, neurite outgrowth and synaptogenesis (Hori et al., 2014, 2020). We hypothesize that AUTS2-l binding at one or more of these intronic enhancers modulates AUTS2-s expression in postmitotic neurons, initiating the transcriptomic cascade required to extend axons and dendrites and develop synapses during late gestation and early postnatal life. Additional data will be required to support this hypothesis. Nevertheless, the finding that AUTS2-l LOF leads to AUTS2-s overexpression is an important clue to the timing, interactions, and the implementation of isoform functions across the developing brain.

The neurodevelopmental pathologies reported here are consistent with the notion that AUTS2 isoforms have distinct and possibly developmental stage-specific functions, including some that are surprising. For example, AUTS2-l is the major isoform expressed in postnatal cerebellum overall and in postnatal Purkinje cells specifically, but the ablation of AUTS2-l resulted only in subtle PC pathologies suggesting developmental delay that was mostly corrected by adulthood. Based on these data, we surmise that the dramatic defects in PC development documented for 16Gso and C-terminal KO mutants (Weisner et al., 2019; Yamashiro et al., 2020) must depend on the activities of AUTS2-s in either the PC themselves or in supporting cells such as differentiating and migrating cerebellar granule cells – possibly during prenatal or early postnatal stages. In contrast, we found that AUTS2-l is essential to hippocampal development, with *Nes-cre,* ex7^-/-^ mice displaying several key hippocampal pathologies documented for C-terminal mutants (Castanza et al., 2021; Hori et al., 2020) and 16Gso mice (Weisner et al., 2019). These shared phenotypes included an overall reduction in hippocampal size and DG gc numbers (Castanza et al., 2021; Li et al., 2022; Weisner et al., 2019), ectopically positioned TBR2+ intermediate progenitors suggesting abnormal migration (Li et al., 2022), and the failed or stalled progression of DG granule neurons from the immature CalR+ to the maturing CalB+ stage (Weisner et al., 2019). In addition to the lower numbers of CalB+ DG neurons, we found that CalB levels were noticeably reduced in the mutant hippocampus; CalB is important for memory consolidation, intracellular calcium level moderation and neuronal protection (Karádi et al., 2012), and the lower levels of CalB expression suggest a functional deficiency in mutant DG gc that did manage to advance to the CalB+ stage. Consistent with this collection of pathological findings, differential gene expression indicated delayed neuron maturation in mutant P0 forebrain together with the reduced expression of genes involved in axon, dendrite and synapse development in mutant hippocampus at P14. The transcriptomic signal was also consistent with the hippocampal pathology associated with C-terminal *Auts2* mutations (Castanza et al., 2021; Hori et al., 2020).

Despite the evidence of delayed DG gc maturation, gene expression data from P14 hippocampus indicated increased levels of neurotransmitter activity, supported by increased hippocampal c-Fos protein expression in P14 mutants and in adults. This suggested a failure of hippocampal inhibitory circuits although hilar mossy neurons, which fail to differentiate or survive in animals homozygous for C-terminal mutations (Castanza et al., 2021), appeared normal in *Nes-cre,* ex7^-/-^animals. First, these data indicate that AUTS2-s is likely required for the differentiation and survival of the HMNs. But furthermore, hippocampal inhibitory circuits were still defective in *Nes-cre*, ex7^-/-^ mice despite HMN survival. The hyper-excitation detected in *Nes-cre,* ex7^-/-^ hippocampus instead was correlated with reduced numbers of AUTS2-expressing PV+ hippocampal interneurons.

Loss or malfunction of hippocampal PV+ interneurons, which control not only hippocampal excitation but also DG gc differentiation during critical postnatal stages (Song et al., 2013), might have been considered as a major driver of AUTS2- associated hippocampal pathology and cognitive memory deficits. However, *Calb1-cre,* ex7^-/-^ animals displayed full NOR deficiencies *without* the loss of PV+ interneurons or the correlated increase in hippocampal c-Fos expression. Furthermore, in *Calb1-cre,* ex7^-/-^ mice, overall DG size and migration of TBR2+ intermediate progenitors were both normal, ruling out a required role for those pathologies in the cognitive defects. Rather, our data point to the delay or block in DG gc maturation - observed in both *Nes-cre* and *Calb1-cre,* ex7^-/-^ animals - as being central to *Auts2-* related cognitive deficits. This delay is likely to reflect the loss of a critical, cell-intrinsic AUTS2-l function within the post-mitotic progenitors, as they shift from the immature CalR+ stage to express AUTS2-l and CalB, begin to extend axons and dendrites, and establish synaptic connections that will persist into adulthood.

The robust association between the *Auts2-*ex7 mutant DEGs and published AUTS2 ChIP peaks suggests hundreds of putative direct AUTS2-l target genes. Many of these putative target genes are associated with neurodevelopmental disease including ASD, ID and related syndromic disorders, and carry out known functions that could explain a host of key behavioral and histopathological mutant phenotypes. Interestingly, although most AUTS2 peaks are positioned near gene promoters (Gao et al., 2014; Liu et al., 2021; Oksenberg et al., 2014), the majority of DEG-associated AUTS2 ChIP peaks were found at distal intronic or intergenic sites including several that have been validated as distal DEG-linked enhancers (Gorkin et al., 2020). These distal AUTS2 binding peaks were associated with a mixture of up-and down-regulated DEGs, including several that were oppositely regulated in P0 FB and P14 HC, suggesting that the effects of AUTS2-l binding could vary with brain region, cell type, and developmental time. The data thus provide some novel and intriguing mechanistic clues about the functions of both major AUTS2 isoform, although many important questions remain to be answered. For example, although *Auts2* mutant brain pathology is most pronounced in hippocampal dentate gyrus and cerebellar Purkinje cells, other brain regions and cell types express *Auts2* at some time in their development, and likely contribute to mutant phenotypes in humans and mice; a full view of behavioral and developmental pathologies will require analysis of isoform function in other brain regions and cell types across the developing brain. Furthermore, evidence suggests that the functions of AUTS2 isoforms are tightly coordinated and probably intertwined, but the biological function of AUTS2-s - the most widely and highly expressed of the two major *Auts2* isoforms – is still mostly a matter of conjecture. Understanding the broad impact of *AUTS2* on neurodevelopment and susceptibility to disease will require a deeper understanding the actions of both AUTS2 isoforms and their interactions within a range of neuronal cell types across the developing brain.

### Data Availability

Sequencing data described in this paper have been submitted to the GEO database under accession number GSE231599.

## Acknowledgements

The authors are grateful to Jennifer Yoo, Derek Sargent, Andrew Look, Manasi Inamdar, and Kian Patton for expert assistance and Kian Patton, Xue Geng, and Stephanie Ceman for comments on the manuscript. We also thank Dr. Danny Reinberg and Dr. James Stafford for providing the *Auts2^tm1.1Dare^* floxed mice and for very helpful discussions and advice. This study was funded by the National Institutes of Mental Health, grant MH114600 (awarded to L.S.) and by the generous support of the Pacific Northwest Research Institute.

---

**Supplementary Figure 1.**
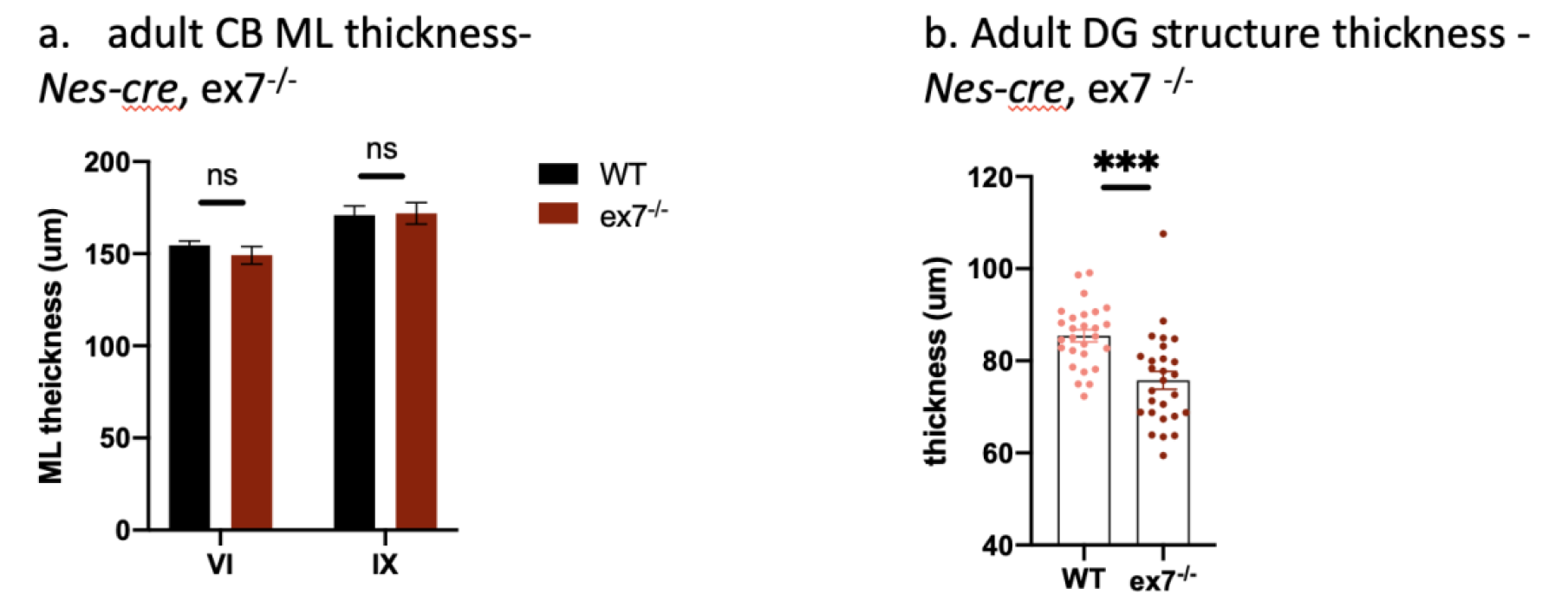
Additional *Nes-cre*, ex7^-/-^ characterizations. **a) Adult cerebellar molecular layer (ML) thickness** was not significantly different between wt and *Nes-cre,* ex7 ^-/-^ mice in lobule VI and IX. **b) Adult DG structure thickness in *Nes-cre*, ex7^-/-^** animals was significantly decreased compared to WT animals. Two-tailed t-test was used to determine statistical significance; ***= p<0.001, ns = not significant.

**Supplementary Figure 2.**
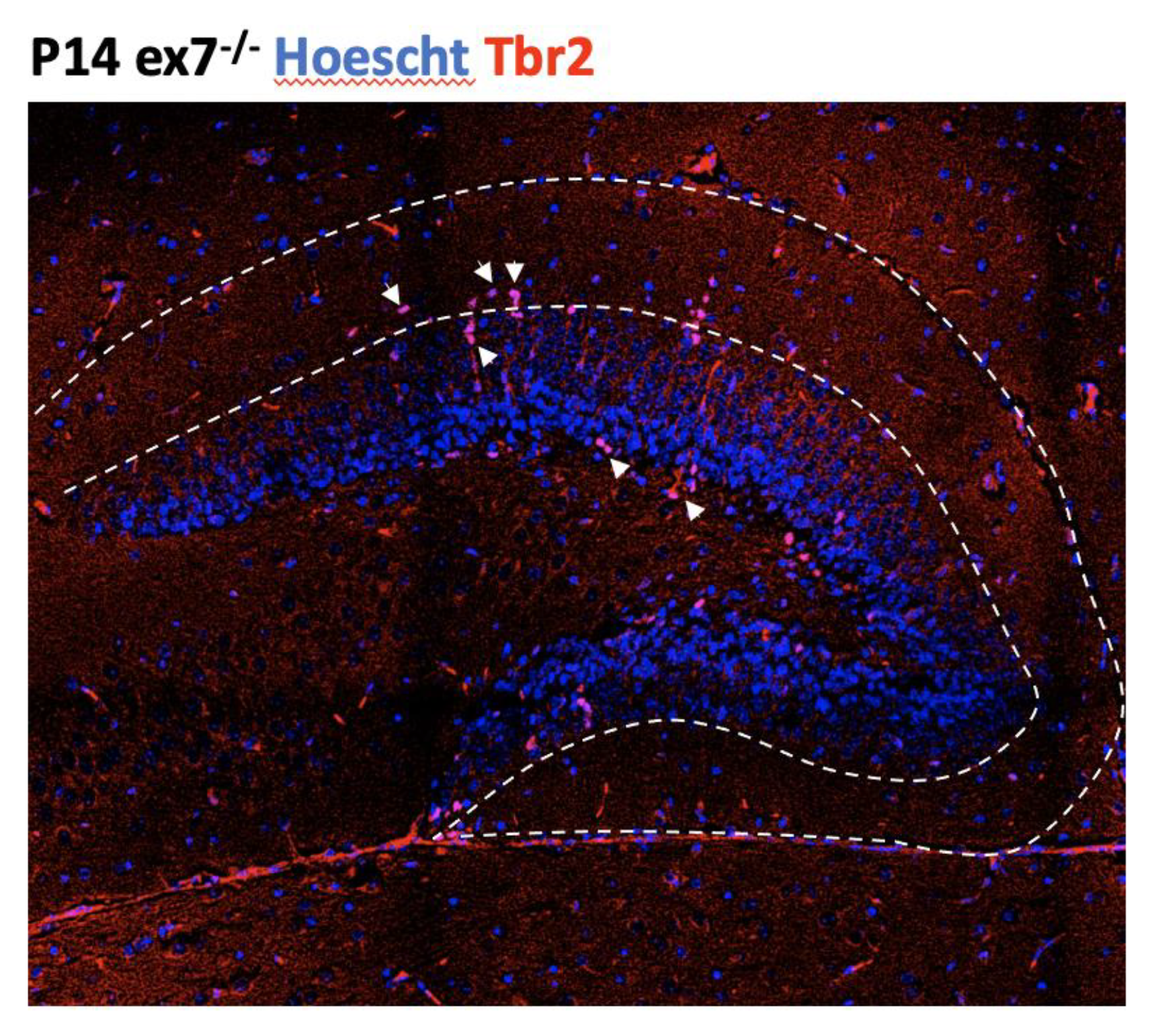
Ectopic location of TBR2+ cells in *Nes-cre*, ex7^-/-^ mutant hippocampus. Representative image of P14 mutant HC with white arrows pointing out TBR2+ cells. TBR2+ cells located between the two white lines represent the “outer” group, whereas cells in the gc layer and in Hilus are included in “Hilus” group.

**Supplementary Figure 3.**
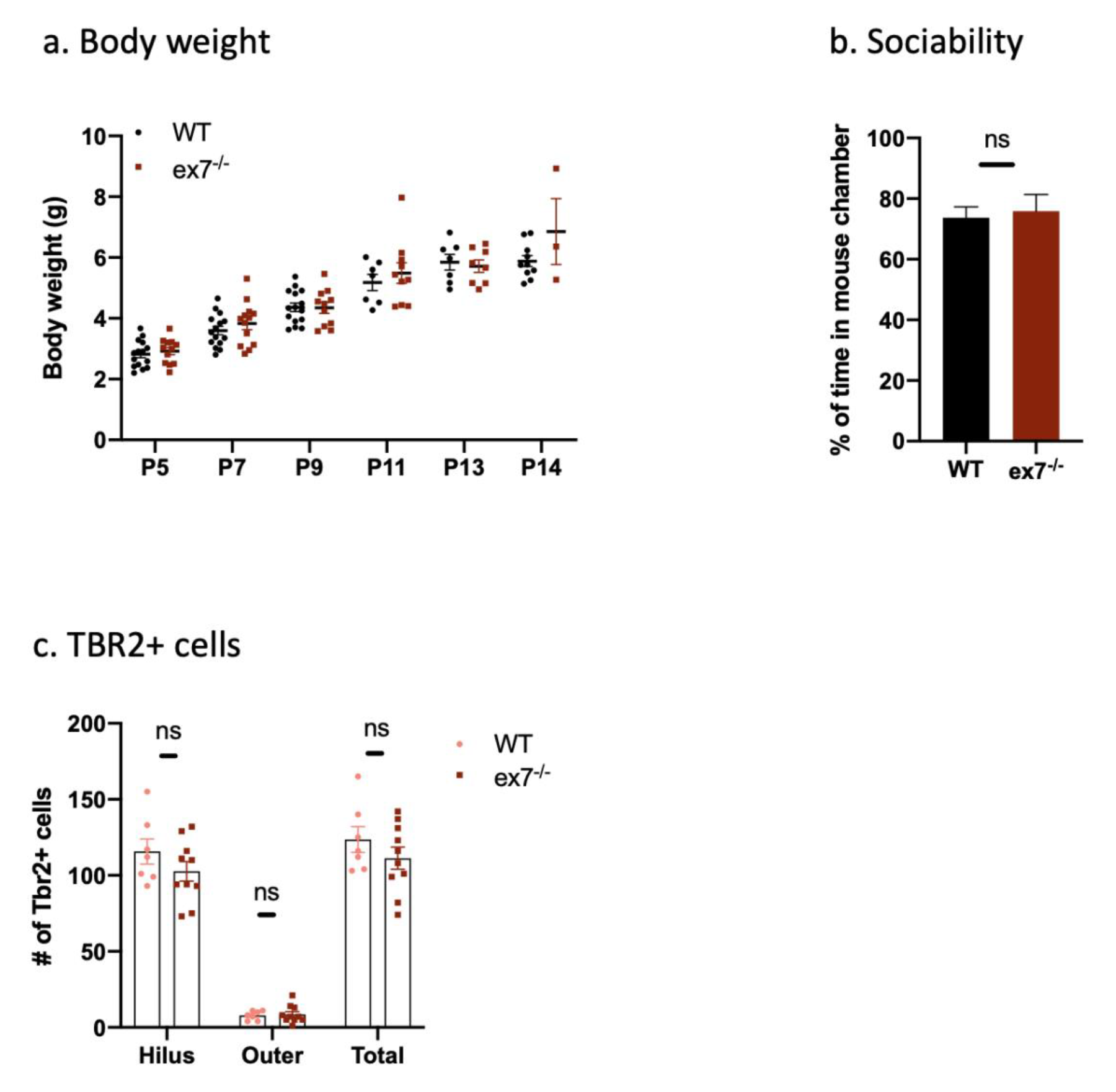
*Calb1-cre*, ex7^-/-^ body weight, sociability, and TBR2+ cell count. *Calb1-cre*, ex7^-/-^ (abbreviated in the figure as ex7^-/-^) a) **Body weight measurements** at different developmental time points. P5, P7, P9, P11, P13, P14, pups were measured for body weight and showed no significant difference to WT animals. WT n= 7-15; *Calb1-cre*, ex7^-/-^ n= 3-13. b) **Sociability.** Panel shows the percentage of time a tested mouse spent in mouse chamber vs. total time spent in mouse chamber plus the toy chamber during second 10 min session of three-chamber test (see method). WT n =8, *Calb1-cre,* ex7^-/-^ n=5 c) **TBR2+ cells in p14 HC.** Neither the total numbers of TBR2+ cells nor their distribution was significantly different in *Calb1-cre*, ex7^-/-^ P14 HC when compared to WT littermates. Two-tailed t-test was used to determine statistical significance; ns = not significant.

**Supplementary Figure 4.**
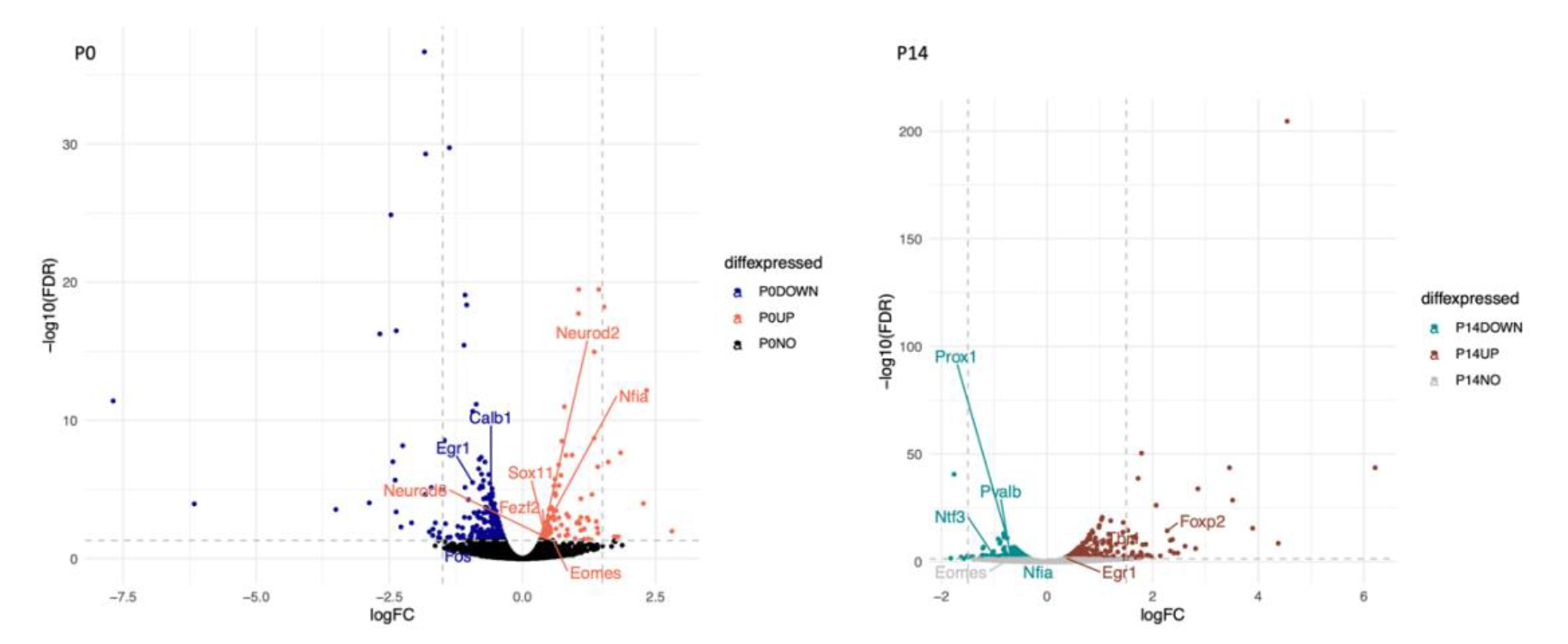
P0 (left) and P14(right) gene expression plot full view. For P0, blue = down-regulated genes, coral = up-regulated genes, black = not differentially expressed. For P14, green = down-regulated genes, brown = up-regulated genes, grey = not differentially expressed. Grey dashed lines represents log FC = −1.5, 1.5, and FDR = 0.05.

